# Conserved atypical cadherin, Fat2, regulates axon terminal organization in the developing *Drosophila* olfactory sensory neurons

**DOI:** 10.1101/2023.07.28.550990

**Authors:** Khanh M. Vien, Qichen Duan, Chun Yeung, Pelin Cayirlioglu Volkan

**Author notes:** Corresponding author 4341 French Family Science Center 124 Science Drive Campus Box 90338, Duke University Durham, NC 27708, USA.

## Abstract

A unique and defining characteristic of the olfactory sensory circuit is its functionally organized topographic map, which requires widely dispersed olfactory sensory neurons with the same identity to converge their axons into one neuropil, a class-specific glomerulus. Understanding the process of how neuronal identity confers circuit organization is a major endeavor in the field of neurobiology due to its intricate connection to neurodegeneration and neuronal dysfunction. In the olfactory system, many aspects of circuit organization, from axon guidance to synaptic matching, are regulated by a variety of cell surface proteins (for example Robo/Slit and Toll receptors). In this paper, we’ve identified a novel atypical cadherin protein, Fat2 (also known as Kugelei), as a regulator of class-specific axon organization. Fat2 is expressed in olfactory receptor neurons (ORNs) and local interneurons (LNs) within the olfactory circuits, but little to no expression is found in projection neurons (PNs). Fat2 expression levels vary in a neuronal class-specific manner and peak during pupal development. In *fat2* null mutants, ORN axon terminals belonging to different ORN classes present with varying phenotypic severity with the highest *fat2* expressing classes being most severely affected. In the most extreme cases, *fat2* mutations lead to ORN degeneration. We then show evidence that suggests Fat2 intracellular domain is necessary for Fat2 function in ORN axon organization. Within the developmental context, Fat2 is required starting at early stages of olfactory circuit development specifically for precise axon retraction to further condense class-specific glomeruli. We’ve also shown that PNs’ and LNs’ expression of Fat2 likely does not contribute to ORN organization ^1^. Lastly, we narrow down potential Fat2 intracellular domain interactors, APC family proteins (Adenomatous polyposis coli) and *dop* (Drop out), that likely orchestrate the cytoskeletal remodeling required for axon retraction during protoglomerular development. Altogether, we provide a foundational understanding of how Fat2 functions in olfactory circuit organization and implicate the critical role of axon retraction during glomerular maturation.

## Introduction

In both insects and mammals, odor detection is highly conserved and depends heavily on diverse classes of olfactory neurons that organize their axons to converge in a class-specific manner within the antennal lobe in flies, or the mammalian central brain’s olfactory bulb ^2^. Given its intricate association with neurodegeneration and neuronal dysfunction, decoding the mechanisms underlying how diverse classes of neurons organize their axons represents a significant pursuit in the field of neurobiology. In flies, olfactory receptor neuron (ORN) classes are defined by the selective co-expression of generally one to three odorant receptor (and/or ionotropic receptor) combinations from a large genomic repertoire of chemosensory genes ^3^. ORNs belonging to the same class converge their axons to a single neuropil, called a glomerulus, and synapse with second-order projection neurons (PNs). This organizational logic is known colloquially as the “one olfactory receptor, one glomerulus” rule. Considering the robustly stereotyped shapes, boundaries, and positionings of the glomerular map, each glomerulus likely requires projections from multiple cell types (presynaptic ORN axons, post-synaptic PN dendrites, and local interneuron (LN) projections) to orderly follow a complex set of instructions. Studies have shown that several aspects of axon behavior, such as axon guidance and axon-axon recognition, can be regulated by a combinatorial code composed of cell surface molecules. Previously, we and others have shown that the disruption of several cell surface molecules, such as Cadherin family proteins, can disrupt class-specific glomerular organization ^4, 5^. Leveraging the field’s detailed understanding of the Drosophila olfactory circuit architecture, we can focus on unraveling the functional relevancy and mechanism of actions behind evolutionarily conserved molecular interactions.

The adult Drosophila olfactory system is an excellent model for studying neurodevelopment because it utilizes mechanisms documented across most multi-cellular neural networks, such as the temporal and spatial regulation of combinations of cell surface molecules to direct the behaviors of neuronal processes. During metamorphosis, adult ORNs are born within the eye-antennal imaginal disc and all ∼1200 ORNs per antenna must extend their axons great distances into the central brain ^6^. Around 16-18 hAPF (hours after pupal formation) ORN axons reach the antennal lobe surface, then choose to join either the ventromedial axon tract or the dorsolateral tract ^7^, and continue along the antennal lobe surface, creating transient exploratory branches and surveying the environment for cues from their target glomerulus ^8^. The mechanisms involved in the stabilization of these transient branches are unknown, and likely involve proteins that link chemoaffinity receptor activity to cytoskeletal regulators. Between 20 hAPF and 40 hAPF, the exploratory axon targets a glomerulus ^9, 10^, begins interacting with PNs and LNs ^11, 12^, and then accentuates proto-glomerulus boundaries by avoiding adjacent glomeruli and further converging with class-specific axons ^4, 13^.The litany of cell surface proteins implicated in olfactory system development, such as Semaphorins, DSCAM, Tenascins/Teneurins and Toll receptors, suggests that ORN class-specific cell surface codes drive this topographic circuit assembly ^9, 11, 13–18^. Given the importance of cell surface combinatorial codes, identifying cell surface proteins functionally required for olfactory circuit assembly will provide the “rosetta stone” needed to translate cell surface signatures into predictable discrete cellular processes. In this study, we focus on the function of a member of the Fat Cadherin family, Fat2, which shows a relatively specific and high expression in the developing olfactory system.

Though there are several examples of cell surface molecules regulating ORN class-specific axon organization, it is still unclear when and how ORN class-specific axon convergence occurs during olfactory circuit development ^4, 15, 16, 19^. The role of Cadherin family proteins has been repeatedly shown to regulate multiple aspects of axon and dendrite behaviors ^20^. Several Cadherin proteins are implicated in olfactory circuit development, including N-cadherin and protocadherins ^4, 21^. However, little is known about the role of the largest Cadherin proteins, one of which is the atypical cadherin Fat2, in the developing olfactory system. Fat cadherins have 34-36 cadherin repeats in their extracellular domain ^4, 22^, which is more than twice the length of Drosophila N-Cadherin, a well-known organizer of protoglomerular organization, which has 17 cadherin repeats. Due to the size of Fat2, a major obstacle to studying the in vivo function of this protein is the difficult and cumbersome molecular biology required to generate functional genetic tools. Recent generation of whole animal null mutants, domain-specific mutants, and GFP-tagged Fat2 direct fusion protein, allows us to functionally characterize this highly conserved protein in a neuronal context ^23–25^.

The appealing aspect of Fat cadherin proteins lies in their remarkable conservation across the animal kingdom. Notably, it has been proposed that the contemporary protocadherin ectodomains may have arisen from the gradual reduction of N-terminal cadherin domains in an ancestral Fat protein, further underscoring the significance of studying these proteins. ^26^. The evolutionary conservation of Fat proteins likely signifies their indispensable role in fundamental biological processes, which when understood could have widespread relevance. Compared to the four Fat cadherins in mammals, Drosophila has two Fat cadherins, Fat and Fat2 (also known as Kugelei), which share striking homology to Fat4 and Fat1/3 in humans, respectively ^27^. It is important to note that the intracellular domains of Drosophila Fat and Fat2 exhibit substantial divergence, characterized by distinct sets of interacting partners that do not overlap. These findings strongly indicate that these two proteins serve distinct functional roles in biological processes. Fat2, and its vertebrate orthologues Fat1 and Fat3, has been shown, using RNA seq, QT-PCR, and in situ hybridization experiments, to be most highly expressed within the developing olfactory bulb and has been associated with several human neuropathologies ^28–30^. The role of Fat2 has been revealed in several mammalian sensory circuits (such as retinal development and cochlear morphogenesis) and Drosophila tissue morphogenesis, yet the function of Fat2 in olfactory circuit development is unknown ^24, 25, 31, 32^

Here we demonstrate that the Drosophila Fat2 protein organizes glomerular architecture by promoting class-specific ORN axon convergence. In fat2 mutants, we observed that many ORN classes displayed fragmented glomeruli with varying severity. By utilizing intersectional genetic approaches, we have identified that Fat2 expression varies across ORN classes during mid-pupal development and is expressed by a unique subpopulation of antennal lobe local interneurons. We have further characterized Fat2 protein localization to axon terminals and provided evidence that the intracellular signaling domain of Fat2 is necessary for its function in glomerular organization. Finally, we present evidence suggesting that Fat2 regulates axon behavior by interacting with cytoskeletal remodeling proteins.

## Results

### Genetic disruptions of Fat2 result in abnormal glomerular organization

Fat3 and Fat1, mammalian orthologues of Drosophila Fat2, are documented to be specifically highly expressed in the olfactory bulbs of developing rats and mice, in addition to other neural circuits ^29, 33, 34^. Consistent with the expression data, several studies show disruptions in Fat3/Fat1 result in a faulty neural organization, in both auditory and visual sensory systems ^35, 36^ in mice. In order to derive the direct impact of Fat2 disruptions on the fly olfactory circuit organization, we restrictively knocked down *fat2* expression using the ORN-specific driver *pebbled-GAL4* (*peb-GAL4*) to drive *fat2* RNAi. To assess the effect of *fat2* knockdown on class-specific glomerular architecture, we visualized one of the largest and most easily identifiable glomeruli innervated by Or47b ORNs using *Or47b-CD2*. We found that compared to *peb-GAL4* only (13% aberrant glomerular morphology, *n*=52) or *UAS-fat2 RNAi* only controls (10%, *n*=54), experimental brains (37%, *n=60*) resulted in Or47b ORNs-innervated VA1v glomerulus becoming morphologically deformed, including glomerular rotation, shape change, and/or fragmentation (Fig 1A). Together this suggests that further studies into Fat2 may elucidate a highly conserved mechanism in organizing developing neural circuits.

**Figure 1.**
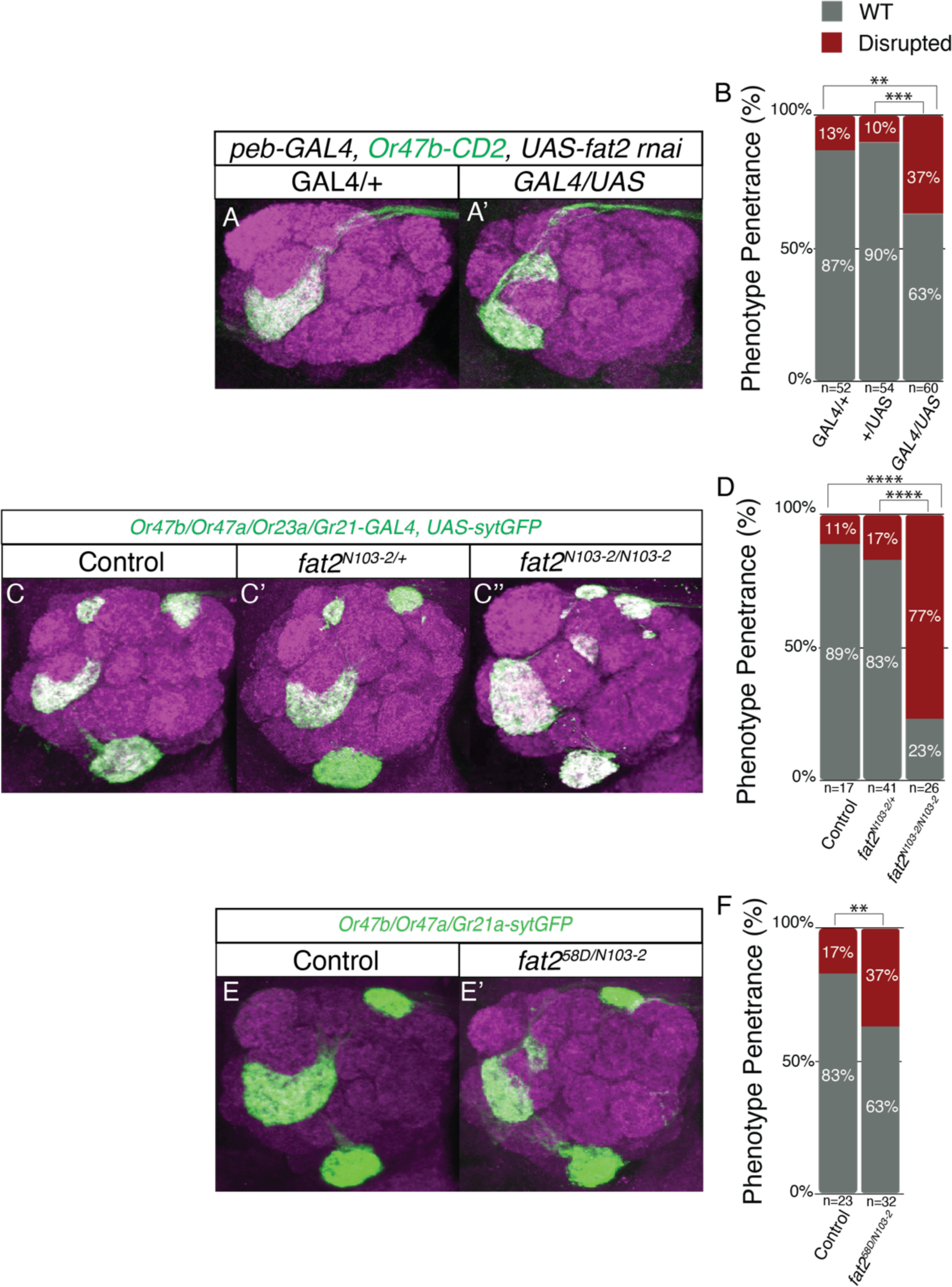
Genetic disruptions in *fat2* breaks stereotype olfactory glomeruli morphology. **(A)** Representative z-stack images of the glomeruli (VA1v) innervated by Or47b axon terminals (green) co-stained with antibody against N-Cadherin (magenta) to visualize antenna lobe structure. ORN-specific (*peb-GAL4*) RNAi knockdown of *fat2* results in a split VA1v glomeruli. **(B)** Quantification of brains with disrupted glomerular morphology for experiment (A) (No *GAL4*) Control (n=52) and (No *UAS*) Control (n=54) are not significantly different. Both controls compared to *peb*>*fat2 RNAi* (n=60) using Fisher’s exact test (**p=.0027,***p=.0004). **(C-C”)** Representative z-stack images of glomeruli innervated by Or47b, Or47a, Or23a, and Gr21a axon terminals (green) co-stained with anti-N-Cadherin (magenta) in Control **(C)**, heterozygotes for *fat2* null (*fat2^N^*^103–2^) **(C’)**, and homozygous *fat2* null whole animal mutants **(C”)***. Or47b, Or47a, Or23a and Gr21a-GAL4, UAS-sytGFP* transgenic flies were used. Homozygous *fat2* null whole animal mutants break apart the continuous neuropil for Or47b, Or47a, Or23a. **(D)** Quantification of brains with disrupted glomerular morphology for Control (n=34), heterozygous for *fat2* null (*fat2^N^*^103–2^) (n=41), and homozygous for *fat2* null (n=26) using Fisher’s exact results in ****p<.0001 **(E)** Trans-heterozygote of two different *fat2* null alleles from separate labs phenocopy homozygous *fat2^N^*^103–2^ mutant phenotype. *Or47b, Or47a, Gr21a* visualized via direct fusion to *sytGFP* and then stained with the same antibodies as (A). **(F)** Using Fisher’s exact test to compare Control (n=23) *fat2^N^*^103–2^ */fat2^58D^*(n=32) using Fisher’s exact test results in significance of **p=.0011

We set out to investigate whether the role Fat2 plays on ORN neuropil organization can also be applied to other glomeruli. To test this, and corroborate our RNAi knockdown results, we visualized four glomeruli (Or47a, Or23a, Or47b, and Gr21a) in whole animal mutants with various *fat2* null alleles. With this paradigm, we were able to compare the morphology of each glomerulus in control brains (11%, *n*=17), homozygous Fat2 null mutant (*fat2^N1^*^03–2^) (77%, *n*=26), and trans-heterozygous mutant for two separately generated *fat2* null alleles (*fat2^N1^*^03–2^/*fat2^58D^*) (37%, *n*=32) (Fig. 1B). *fat2^N^*^103–2^ is a null allele containing a premature stop codon at position 3718 ^23^, and *fat2^58D^* is a null allele that contains a deletion for the amino acids 1-687 including the start codon ^37^. In these whole animal mutants, we observed three glomeruli targeted by Or47a, Or23a, and Or47b ORNs exhibited an organizational disruption. Supporting our previous data, we found that whole animal mutants phenocopy ORN-specific *fat2* RNAi knockdown animals in VA1v glomerulus. We interpreted the presence of ectopic glomeruli and ectopic axon projection as possibly a result of a loss of class-specific axon aggregation. Heterozygous Fat2 null mutants did not exhibit any glomerular disruptions (Fig 1C’).

To summarize, both ORN-specific RNAi knockdown of *fat2*, as well as whole animal mutants for homozygous and transallelic *fat2* null alleles results in disruptions in the organization of multiple glomeruli across the antenna lobe. Indeed, consistent with our finding, a previous study also reported the disrupted Drosophila olfactory circuit architecture in pan-neuronal knockdown of *fat2* ^38^. This suggests that Fat2 plays a functional role in olfactory circuit organization.

### Fat2 expression changes throughout olfactory system development

One possible mechanism by which Fat2 regulates glomerular organization is by establishing an expression level signature unique to each ORN class. To get a brief idea of which ORN class(es) express Fat2, we analyzed published single-cell RNA-seq data to produce a list of Fat2-expressing ORN classes (Fig 2A)^30^. Based on our analysis, Fat2 seems to be broadly expressed across ORN classes (Fig 2A). Strikingly, Fat2 is much more highly expressed at 24 hAPF (hours after pupa formation) and 48 hAPF than in the adult ORNs. This expression pattern supports our previous evidence that Fat2 functions during olfactory circuit development to organize class-specific synaptic neuropils.

**Figure 2.**
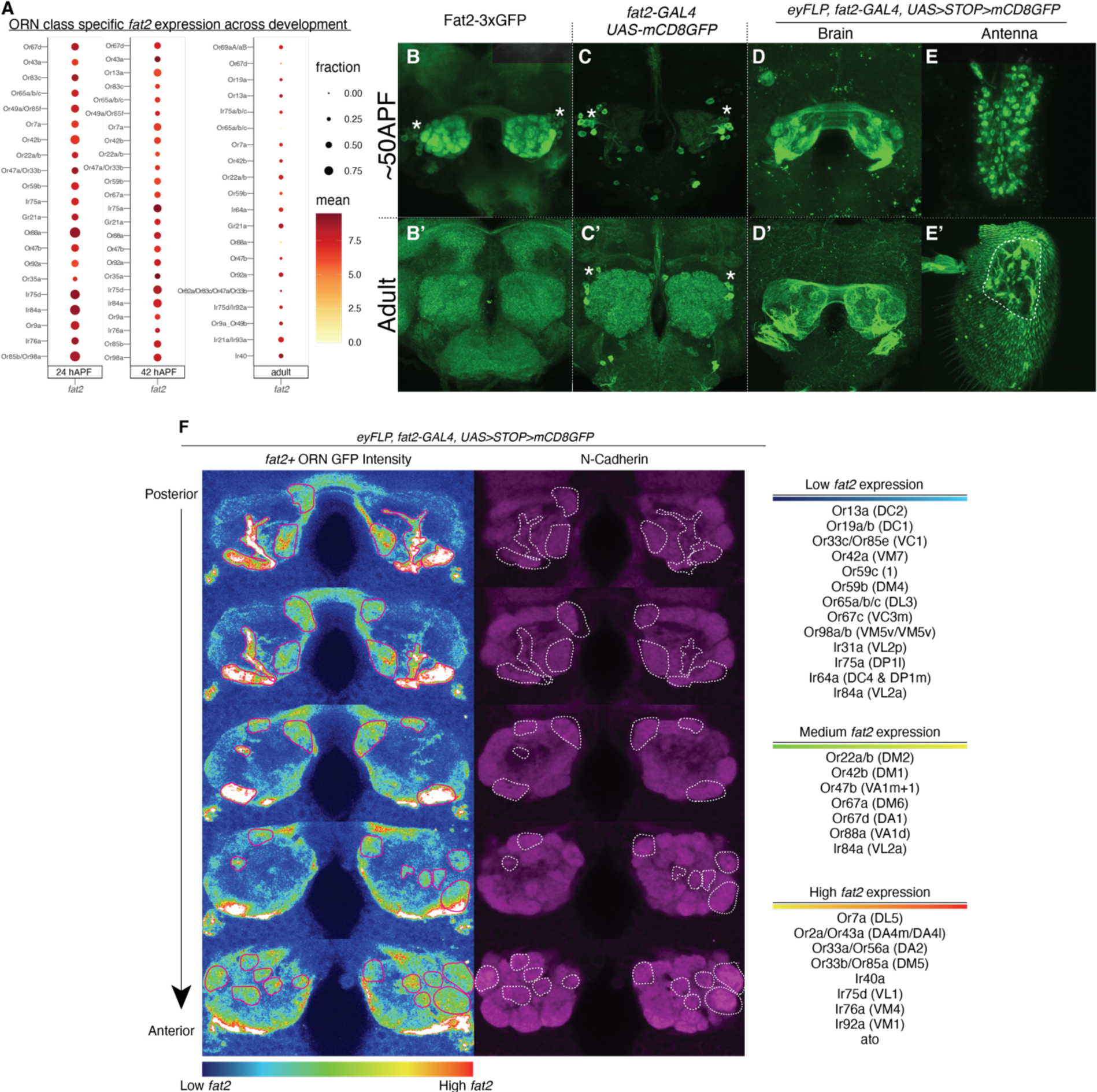
Pupal olfactory system shows developmentally distinct *fat2* expression and Fat2 protein sub-cellular localization. **(A)** Dot plot summary of *fat2* expression per select ORN class at 24 hAPF, 48 hAPF, and adult using previously published single cell RNA-seq datasets. **(B-E, B’-E’)** 50 hAPF pupal brains and adult brains stained with anti-GFP for Fat2-3xGFP (fat2 directly fused to GFP), *fat2*-*GAL4* driven *UAS-mCD8GFP*, and intersectional labeling of *fat2*-positive ORNs using *eyFLP*, *fat2-GAL4, UAS>STOP>mCD8GFP.* Antennal images also stained with anti-GFP for *eyFLP*, *fat2-GAL4, UAS>STOP>mCD8GFP* to visualize *fat2*-positive ORN cell bodies. Cell bodies of unknown identity adjacent to the AL are also positive for *fat2* expression, marked with white asterisks. (F) *eyFLP*, *fat2-GAL4, UAS>STOP>mCD8GFP* pupa at 70 hAPF. Heatmap representing GFP fluorescence intensity and N-Cadherin antibody stain in magenta to identify glomeruli position. Stereotypical location and shape of each glomerulus are visually assessed and assigned an ORN class based on olfactory maps described in the literature^2, 3, 39^

Unfortunately, this antennal single-cell RNA-seq dataset captures only a limited portion of all ORN classes. Therefore, to uncover a more complete *fat2* expression profile, we chose an genetic intersectional approach, using *ey-FLP* and *fat2-GAL4* to restrict the expression of reporter mCD8GFP in *fat2*-positive ORNs specifically. We analyzed GFP expression during olfactory system development to visualize the developmental dynamics and heterogeneity of *fat2* expression across ORN populations. We found that Fat2 is not expressed by ORNs until after 24 hAPF (Supp. Fig 1). Fat2 expression within the antennal lobe begins between 30-40 hAPF. We noticed variable levels of green fluorescence across glomeruli suggesting differential *fat2* expression levels across ORN classes (Fig 2 B-D’, and F). In adults, we found that ORNs belonging to Ir40a, Ir75d, and Ir76b classes and their respective glomeruli showed the highest levels of Fat2 expression ^39^ (Fig 2F). These results reaffirm the list of *fat2*-expressing ORN classes in the single-cell RNA-seq data (Fig 2A). Even though inter-glomerular fluorescence signal variation seems to be maintained throughout pupal development, overall antennal lobe fluorescence is highest between 42 hAPF to 48 hAPF (Supp. Fig 1). Aside from glomeruli targeted by Ir (ionotropic receptor) ORNs, *fat2* expression was also higher for glomeruli targeted by Or19a, Or13a, and Or56a ORNs. The lowest levels of *fat2* expression were observed for glomeruli targeted by Ir64a and Ir84a ORNs. To confirm that the glomerular fluorescence signal is coming from ORN projections and not from other antennal lobe neurons, we analyzed the GFP fluorescence in the antennae, where ORN cell bodies reside (Fig 2 E-E’). We found an abundance of *fat2-*positive ORN cell bodies throughout the mid-pupal antennae (Fig 2E), suggesting that the majority of ORNs express Fat2 during pupal development. Yet as the fly matures, high Fat2 expression becomes restricted to ORN cell bodies originating from the sacculus in the adults (Fig 2E’). These results suggest that 1) f*at2* is expressed in an ORN class-specific manner and 2) *fat2* expression begins around 30 hAPF coinciding with the beginning of protoglomerular formation.

If Fat2 is expressed in ORNs to regulate the class-specific organization of axons and axon terminals, we would predict Fat2 to localize along ORN axons and synapses as opposed to ORN cell bodies. To probe Fat2 protein sub-cellular localization, we leveraged a CRISPR-generated transgenic Drosophila line with three copies of GFP tagged to the C-terminus of Fat2 (Fat2-3xGFP) ^25^. We found GFP fluorescence in the brain was much higher during pupal stages compared to that in adults, which is consistent with data from single-cell RNA-seq and *fat2-GAL4* expression patterns (Fig 2B-B’). Further supporting the functional role of Fat2 in organizing ORN axons, we found that Fat2-3xGFP fluorescence was predominantly confined to the antennal lobe and commissural tract, which indicates localization along ORN axon projections and terminals (Fig 2B). On the other hand, we did not see any fluorescence in either pupal or adult ORN cell bodies on the antennae (Supp. Fig 3), suggesting that Fat2 protein is indeed primarily localized at the neuronal projections within the antennal lobe during development.

**Figure 3.**
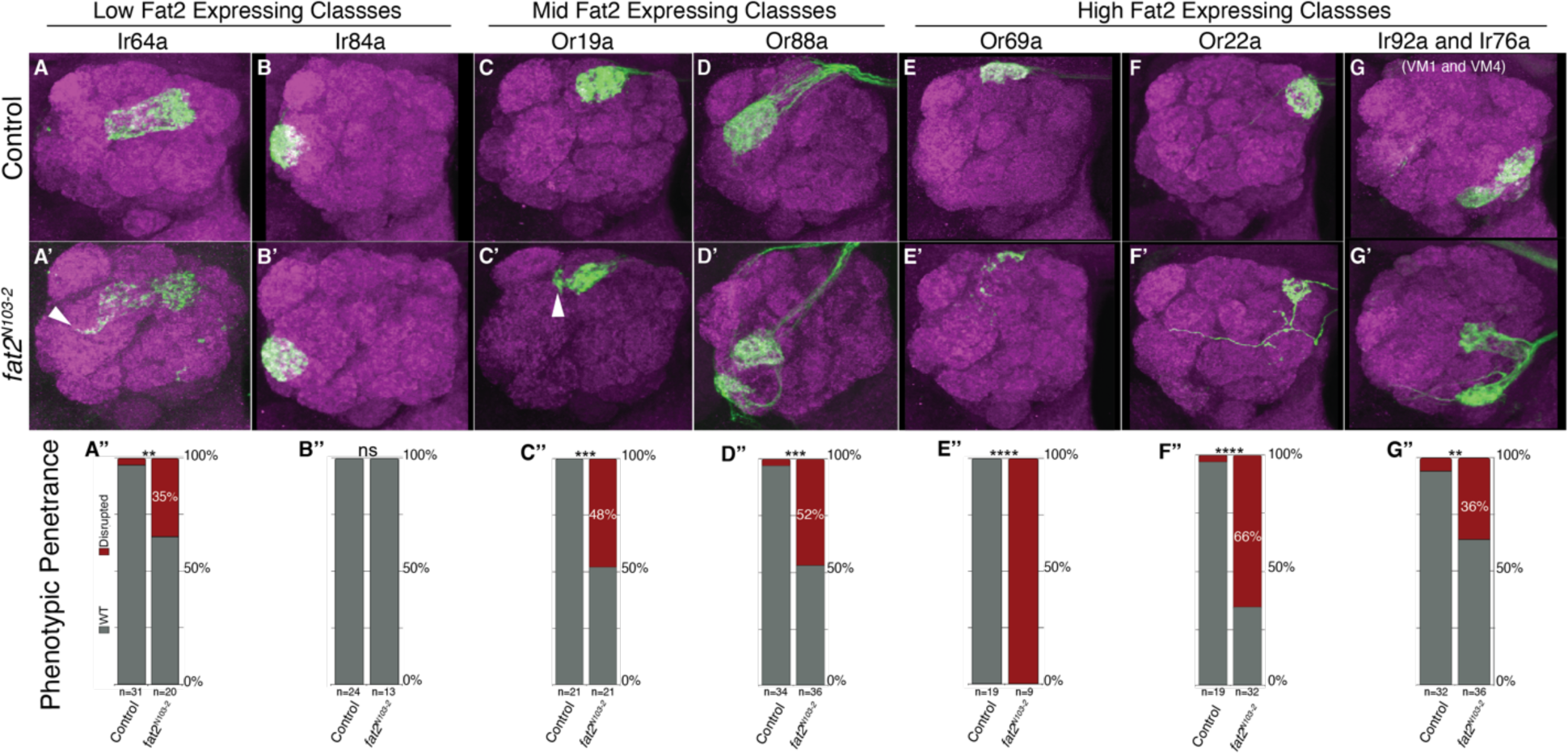
Class-specific *fat2* expression levels predict severity of *fat2* null mutant phenotypes. (A-G, A’-G’) Representative z-stack composite images of single antenna lobes visualizing two low *fat2* expressing ORN classes (Ir64a and Ir84a), two mid *fat2* expressing classes (Or19a and Or88a), and three high *fat2* expressing classes (Or69a, Or22a, Ir92a, and Ir76a). Control and *fat2^N^*^103–2^ homozygotes are stained with anti-GFP (green) to show axon terminals and anti-N-Cadherin (magenta) for AL gross anatomy. Each ORN class is labeled using *OrX-GAL4* (or *72OK-GAL4* for Ir92a and Ir76a) and *UAS-SytGFP*. Aberrations in glomerular structure (emphasized with white arrows) seen in mutant alleles but rarely, if ever, appear in control brains. **(A”-G”)** Bar graphs quantifying phenotypic penetrance in Control and fat2 null homozygouse (A-G’). Fisher’s exact test performed on quantifications (A’) 35% disrupted phenotype in *fat2^N^*^103–2^ mutants, p=.004 (B”) 0%, not significant (C”) 48%, p=.0005 (D”) 52%, p<.0001(E”) 100%, p<.0001 (F”) 66%, p<.0001 (G”) 36%, p=.007

Collectively our results indicate that Fat2 expression is localized to ORN axons and synaptic terminals, shows class-specificity in its expression levels, and peaks between 30 hAPF to 75 hAPF, suggesting Fat2 likely plays a role in establishing the functional topography of the glomerular map during pupal development.

### Fat2 expression level predicts class-specific glomerular phenotypes in *fat2* mutants

We then set out to test how the differential expression levels of *fat2* across ORN populations contribute to the class-specific glomerular organization. Using the results from the intersectional labeling approach and the known glomerular map, we were able to identify and categorize ORN classes into high and low *fat2* expressing classes (Fig 2F) ^2, 3, 40^. Then using this list and the available transgenic tools to label each ORN class, we analyzed the glomerular organization in wild type and *fat2* null (*fat2^N^*^103–2^) whole animal mutants. To determine the effect of *fat2* mutations on ORN classes expressing high levels of Fat2, we analyzed D, DM2, VM4, and VM1 glomerular morphology targeted by Or69a (100% disrupted phenotype, *n=9*), Or22a (66%, *n=32*), Ir76a and Ir92a (36%, *n=36*) ORNs (Fig 3E-G). For classes that express medium levels of Fat2, we analyzed DC1 and VA1d glomeruli innervated by Or19a (48%, *n=21*) and Or88a (52%, *n=36*) ORNs (Fig 3C-D). And lastly to assess glomeruli expressing low levels of Fat2, we analyzed DC4/DP1m and VL2a glomerular morphology targeted by Ir64a (35%, *n=21*) and Ir84a (0%, *n=13*) ORNs (Fig 3A-B’). The most common morphological perturbation we found across ORN classes that show a phenotype is a fragmentation or discontinuity of what should be a single glomerulus. Glomerular phenotypes in *fat2* null mutants vary in penetrance in a class-dependent manner where high Fat2 expressing classes were the most susceptible to *fat2* genetic disruptions, whereas low Fat2 expressing classes were the most resistant to Fat2 disruption. High Fat2-expressing classes also presented with more distinctly fractured neuropils in *fat2* mutants in contrast to the mild structural distortions seen in low Fat2-expressing class-specific glomeruli. In addition to disruptions in glomerular morphology, we found that glomerulus D innervated by Or69a ORNs completely disappeared in *fat2* mutants.

This is likely due to decreasing ORN cell number since 7+-day-old flies homozygous for *fat2* null alleles exhibit antennal degeneration (Supp. Fig 2). The variation in glomerular phenotypes suggests that ORN classes show differential requirement for Fat2 function, likely based on expression levels.

In summary, due to ORN class-specific differences in the level of Fat2 expression, we observe variable penetrance and expressivity of the glomerular disruption phenotypes for ORN classes analyzed. Interestingly, Or69a ORNs, which are among the ORN classes with the highest levels of *fat2* expression, are one of the most susceptible classes to loss of *fat2* function. Meanwhile, Ir84a ORNs, which are among the classes with the lowest levels of *fat2*, appears wild type in *fat2* mutants.

### Fat2 intracellular domain is necessary for appropriate glomerular organization

Fat2 is predicted to function as a cell adhesion molecule, which upon trans interactions across neuronal membranes lead to cytoskeletal remodeling to support axonal fasciculation and axon terminal morphology ^41^. To test whether the intracellular domain is required for Fat2 function in ORNs, we analyzed *fat2* mutants that encode a truncated Fat2 protein with the intracellular domain deleted (*fat2^ΔICD^*). *fat2^ΔICD^* mutants effectively phenocopy the Fat2 null mutants for Or47b, Or47a, and Or23a glomeruli (75% disrupted brains, n=36) (Fig1, Fig 4A and 4A’). This suggests signaling from Fat2 intracellular domain is required for Fat2 function during glomerular organization. We further tested whether the intracellular domain of Fat2 was responsible for the ORN class-specific phenotypes (Fig 4B-D’). Again, ORN classes with low Fat2 expression experienced little to no disruption in glomerular organization, whereas ORN classes with high Fat2 expression, exhibited strong disruptions. Surprisingly, heterozygous *fat2^ΔICD^* mutants also display disruptions in glomerular morphology, not seen in *fat2* null heterozygous animals (*fat2^ΔICD^* heterozygous 62% phenotypic penetrance, *fat2* null heterozygous 17% phenotypic penetrance) (Fig 1C’ and Fig 4A), suggesting possible dominant negative effects from Fat2^ΔICD^ protein. In summary, Fat2 intracellular domain plays a critical role in Fat2 function as an organizer of ORN class-specific glomerular patterning.

**Figure 4.**
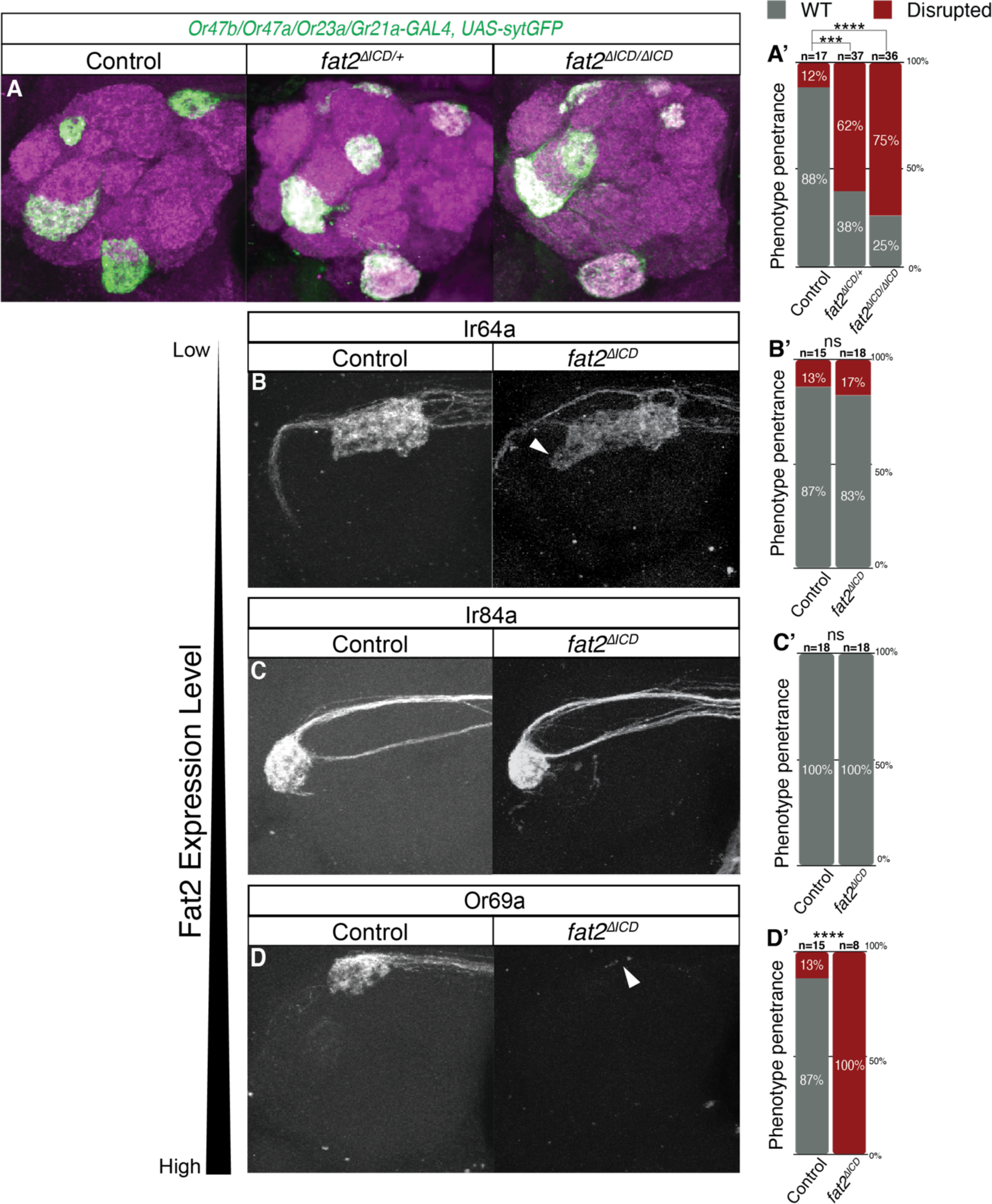
Fat2 intracellular domain mutant, with intact extracellular domain, phenocopies *fat2* null mutant glomerular phenotype in a class-specific manner. **(A-D)** Representative z-stack composite images of single antenna lobes. Only panel (A) is stained with anti-GFP and anti-N-Cadherin. All other images were unstained. In (B-D) each ORN class is labeled using the same *OrX-GAL4* as (Figure 3) but instead drives the expression of *10xUAS-RFP*. *fat2^ΔICD^* brains were from flies homozygous for the *fat2^ΔICD^* allele. **(A’-D’)** Bar graph quantification of brains with glomerular disruptions (red bar). Only bar graphs in (A’) are significantly different as well as bar graphs in (D’). Fisher exact test for (A’) and (D’), separately, resulted in ****p<.0001 and ***p=.0009.

### Fat2 regulates ORN axon retraction during early protoglomerular development

Having established that Fat2 intracellular domain plays a role in axon organization, it is likely that it propagates a signal transduction cascade to govern axon behavior. During olfactory circuit development ORN axons must perform several behaviors, including axon guidance towards the antennal lobe, axon targeting to the appropriate region within the antennal lobe, class-specific bundling to form proto-glomeruli, and lastly establishing synapses with second-order neurons to lead to the formation of mature glomeruli ^8^. To assess the impact of Fat2 on axon behavior during glomerular development, we first sought to identify the earliest developmental time point when we can visualize aberrant glomerular organization. Given that Fat2 expression begins at around 30 hAPF, we analyzed two time points after the onset of Fat2 expression. To perform the developmental analysis, we analyzed three glomeruli targeted by Or47a, Or47b, and Or23a ORNs at 48 hAPF and 72 hAPF. We found that at 48 hAPF, glomerular morphology was already disrupted for the three ORN classes where split glomeruli were visible (Fig 5A). Additionally, the disrupted glomerular morphology is maintained at 72 hAPF and into adulthood. This suggests that glomerular morphology is likely stabilized by 48 hAPF and indicates that the phenotypes likely arose in earlier developmental time points.

**Figure 5.**
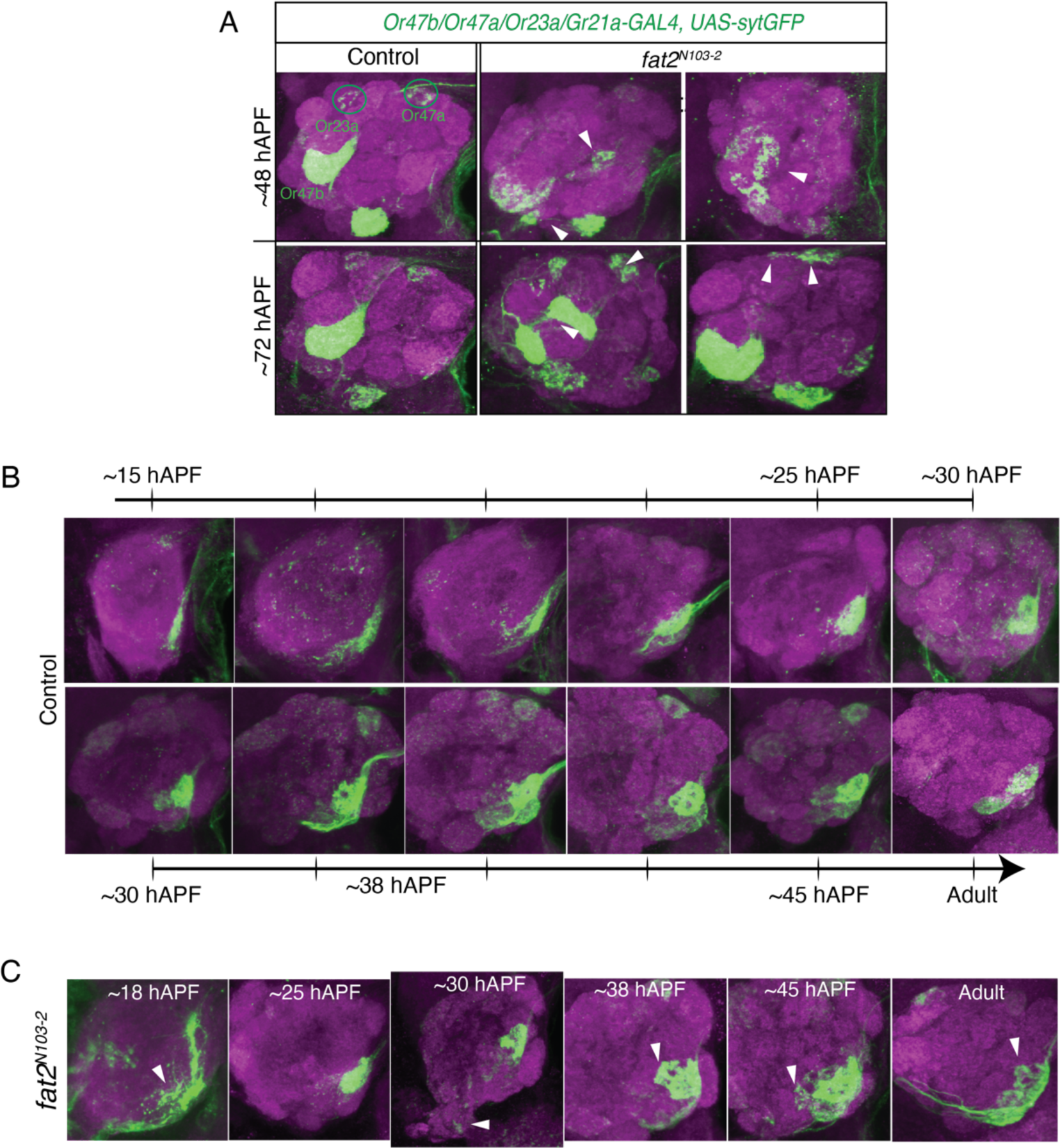
*fat2* mutant phenotype is apparent during protoglomerular development between 18 hAPF to 45 hAPF. **(A)** Representative images of control and *fat2^N^*^103–2^ show three glomeruli corresponding to Or23a, Or47a, Or47b at 48 hAPF (hours after pupal formation) and 72 hAPF, which corresponds to 50% and 75% pupal development respectively. Brains were stained with anti-GFP to visualize ORNs and with anti-N-Cadherin to visualize AL gross anatomy (magenta). White arrows point to disrupted glomerular morphology. **(B)** Representative images of glomeruli labeled by *72OK-GAL4* driver in control brains between 15 hAPF and 45 hAPF. Developmental trajectory of *72OK-GAL4* axons shows two distinct extension phases, one from 15 hAPF to 25 hAPF and the second from 30 hAPF to 45 hAPF. By 45 hAPF, glomerular morphology closely resembles adult glomerular morphology. **(C)** Representative images of *fat2^N^*^103–2^ pupal brains between 18 hAPF and 45 hAPF visualizing axons labeled by *72OK-GAL4* (green) and N-Cadherin (magenta). White arrow points to aberrant axonal projections.

Because most *Or* genes are not expressed until 40 hAPF, we used an early developmental marker *72OK-GAL4* which strongly labels two Fat2-positive glomeruli (VM1 and VM4) with glomerular defects in *fat2* mutants (Fig. 3E-E’). *72OK-GAL4* driven *UAS-sytGFP* expression labels Ir92a (innervates VM1 glomerulus) and Ir76a (innervates VM4 glomerulus) ORNs within the first 15 hAPF ^4, 40, 42, 43^. This allows us to visualize axon behaviors from the earliest points of pupal stages through adulthood. We found that in control brains from 15 hAPF to about 22 hAPF the axons extend dorsal-laterally, then around 25 hAPF the axons retract and condense into the spherical boundaries of VM1 protoglomerulus (Fig 5B). Around 30 hAPF there is a second phase of axon extension towards VM4 which also results in an irregularly shaped VM1 glomerulus. VM1 and VM4 glomerular morphology seems to stabilize after a second axon retraction and neuropil condensation phase between 40 hAPF and 50 hAPF, after which glomerular morphology is stable until adulthood. In contrast to control brains, we found that in *fat2* null mutants the axons did not seem to undergo the second phase of axon retraction and neuropil condensation (Fig 5C). The first phase of axon retraction around 25 hAPF is successful; however, the second phase of retraction around 40 hAPF is often lost. This results in mature VM1 and VM4 glomeruli splitting and exhibiting ectopic target glomeruli. Interestingly, the second phase of retraction coincides with the onset of Fat2 expression within the antennal lobes.

In conclusion, *fat2* mutants disrupt axon behaviors during protoglomerular formation in early pupal development. We found that in *fat2* mutants, ORN axons lose the ability to retract and condense into ORN class-specific protoglomerular fields.

### Abnormal glomerular organization in Fat2 mutant is a result of ORN-ORN interactions

Given that previous research found that Fat2 is able to homodimerize in trans, we set out to identify if any other cells projecting into the antennal lobe also expressed Fat2 and whether their expression is required for glomerular organization ^44^. In the brains of transgenic animals expressing Fat2-3xGFP direct fusion proteins, we did find a population of Fat2-positive cells with cell bodies located adjacent to the antennal lobe and neurites projecting into the antennal lobes (Fig 2B,C). There are two known cell populations with cell bodies adjacent to the antennal lobe, projection neurons (PNs) and local interneurons (LNs) ^43^. To ascertain the neuronal identity of the Fat2-positive cell bodies, we conducted co-labeling experiments with specific markers for both PNs and LNs. By assessing the extent of cellular overlap, we aimed to identify the neuron type(s) associated with the Fat2-positive population. We first labeled a majority of PNs with *GH146-GAL4* driven expression of *10xUAS-RFP* ^45^ in the background of endogenous GFP-tagged Fat2 (Fig 6A). We saw minimal to no overlap between PNs and the Fat2-3xGFP signal (Fig 6A). Additionally, intersectional genetic systems that selectively label *fat2-*positive PNs revealed only one to three PN somas per antennal lobe innervating DM2 and VM2 glomeruli, which are innervated by Or22a and Or43b ORNs respectively (Supp. Fig 3). We also used *fat2-GAL4* to drive the expression of DenMark, a dendrite-specific marker that would highlight any glomeruli innervated by *fat2-GAL4-*positive PNs, if any (Supp. Fig 3) ^46^. We again found that *fat2*-positive dendrites were much fainter compared to the *GH146-GAL4-*labeling dendrites (Supp. Fig 3). Together, we used multiple genetic strategies to show that few PNs express Fat2 and glomerular Fat2 signal in the antennal lobes can be attributed to Fat2 localizing to ORN axon terminals, not PN dendrites.

**Figure 6.**
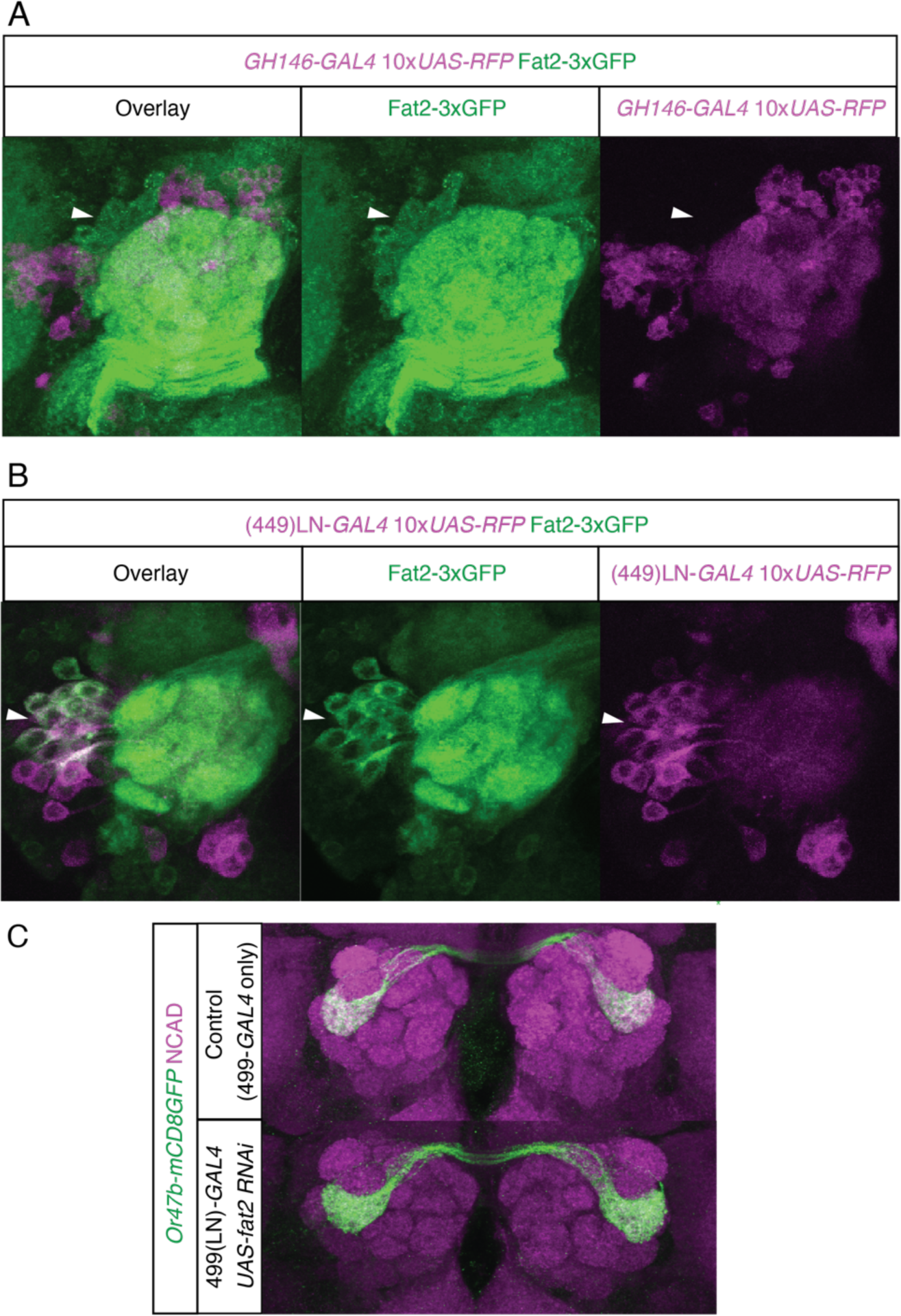
Local interneuron expression of *fat2* does not contribute to ORN glomerular organization. **(A)** Representative unstained z-stack image of both AL where *GH146-GAL4*-labeled PNs in magenta and Fat2-3xGFP fusion protein in green. White arrows represent Fat2+ cell bodies that do not colocalize with PN cell bodies. **(B)** Similar to panel A, Fat2-3xGFP fusion protein is colorized green and *499-GAL4* positive LNs are colorized magenta. White arrows represent Fat2+ cell bodies that colocalize with LN cell bodies. **(C)** Representative z-stack composite images of control AL and LN-specific knockdown of *fat2 (499-GAL4, UAS-fat2 RNAi)* stained with anti-N-Cadherin (magenta) and Or47b axons projecting into corresponding glomeruli (Green) stained with anti-GFP (*Or47b-GAL4, UAS-mCD8GFP*).

**Figure 7.**
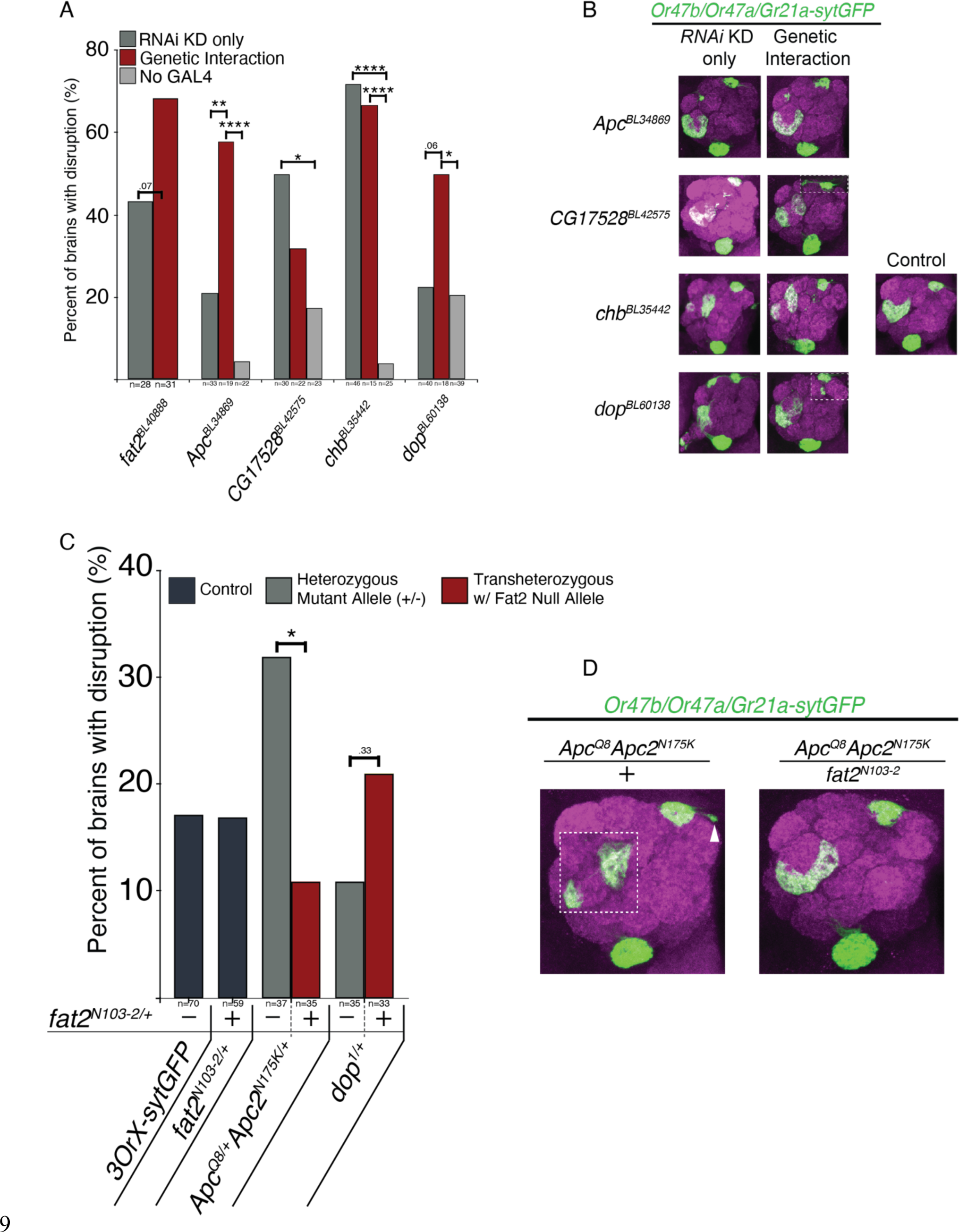
Testing genetic interactions with *fat2* by knocking down putative Fat3 intracellular domain interactors in sensitized *fat2* null heterozygous background. **(A)** Bar graph quantifying percentage of brains with disrupted glomerular morphology (in B) to test genetic interactions between *fat2* and the respective genes. Dark grey bar represents ORN-specific RNAi knockdown only for the denoted gene. Red bar represents RNAi knockdown of denoted gene in *fat2* null heterozygous background. Light grey bar represents control group with no *GAL4*, just *UAS-RNAi* in *fat2* null heterozygous background. *fat2* RNAi knockdown with and without one copy of *fat2^N^*^103–2^ used as positive control (p-value calculated using fisher’s exact, p=.0695). *p<.05, **p<.01, ****p<.0001. **(B)** Representative confocal z-stack composite images of the disrupted glomeruli morphology seen in each group. Brains were stained with anti-N-Cadherin in magenta to visualize antennal lobe anatomy, and anti-GFP in green. Three ORN classes are labeled: Or47a, Or47b, and Gr21a. Dotted white boxes show overlay from another brain with representative disruptions to Or47a (or Gr21a) innervated glomerulus. **(C)** Bar graph quantifying whole animal transheterozygote analysis of one copy of *dop* or *Apc/Apc2* null allele with and without one copy of *fat2^N^*^103–2^. Black bars represent two control conditions, gray bar represent *dop* or *Apc/Apc2* null heterozygotes, red bar represent transheterozygotes disrupting one copy of the gene of interest (*dop* and *Apc/Apc*) and one copy of *fat2*. Differences between red bar and gray bar suggest genetic interaction between the gene of interest and *fat2*. **(D)** Representative images of glomerular disruptions found in specified genotypes. Dotted white boxes show overlay from another brain with representative disruptions to Or47b ORN innervated glomerulus, and white arrow points to fracturing glomerulus innervated by Or47a ORNs.

LNs also have cell bodies adjacent to the antennal lobe and project neuronal processes into the antennal lobe glomeruli. To positionally compare the *fat2*-positive somas to the cell bodies of several LNs subclasses ^47^, we again co-labeled LN subclasses using LN-specific GAL4s driving *UAS-RFP* together with Fat2-3xGFP. We found only one LN subclass labeled by *499-GAL4* that overlapped with Fat2-3xGFP-positive cell bodies (Fig 6B). Given the proximity between LN axons and ORN axons, it is likely that Fat2 could regulate glomerular organization through mediating interactions between developing LN projection terminals and ORN axon terminals. To investigate this possibility, we knocked down *fat2* specifically in *499-GAL4-* positive LNs. We found no noticeable disruption to the axon organization for Or47b ORNs (Fig 6C), nor to LN axon organization (Supp Fig 4). This suggests that local interneuron’s expression of Fat2 does not play a role in establishing glomerular morphology or ORN axon organization.

Together, our results revealed that outside of ORNs, Fat2 is also expressed in LN classes but not PNs within olfactory circuits. Yet LN expression of Fat2 is not required for its function in glomerular organization. Given that 1) Fat2 is predominantly expressed by ORNs and LNs and 2) only ORN-specific knockdown of *fat2* results in glomerular disruptions, ORN axon convergence is likely mediated by Fat2 localization at ORN terminals to drive ORN-ORN interactions.

### Fat2 intracellular domain orchestrates complex interactions with cytoskeletal regulators

To determine how ORN-ORN interactions can turn into Fat2-induced intracellular signaling that organizes ORN glomerular terminals, we screened proteins that interact with Fat2 intracellular domain and play roles in regulating glomerular organization. To narrow down our target list, we used a recently generated dataset of mammalian Fat3-ICD interactors identified using protein pulldowns and mass spectrometry ^48^. Among the 103 proteins in the dataset, we selected the top 10 potential interactors that had Drosophila orthologues with available RNAi genetic reagents to assess glomerular organization in knockdown conditions. *peb-GAL4* driven *UAS-RNAi* knockdown identified four of these genes (*Apc, CG17528* (*zyg8*)*, chb, dop*) phenocopy glomerular organization defects found in *fat2* mutants (Supp Table 1). We also confirmed the phenotype was not caused by the background effect associated with the *UAS-RNAi* transgene docking site (Supp Fig 5), which may influence the ORN glomerular organization based on our previous finding ^49^.

Based on the literature *CG17528* and *Apc* play important roles in regulating actin and microtubule cytoskeleton, while *dop* encodes a kinase that regulates protein localization and transport ^50–64^. If Fat2 intracellular domain indeed interacts with these proteins, we imagine that Fat2 gathers the appropriate cytoskeletal effectors to facilitate axon behavior. We further investigated whether any of these genes genetically interacted with *fat2* by asking if reducing the dose of *fat2* by half in the ORN-specific RNAi knockdown background enhanced or suppressed phenotypes observed (Fig 8A-B). This secondary screen identified *dop* and *Apc* RNAi knockdown glomerular phenotypes to be enhanced if they also carried a copy of *fat2* mutation (Fig 8A). On the other hand, *fat2* mutation mildly suppressed RNAi phenotypes observed in *zyg8/CG17528* knockdowns (Fig 8A-B). In addition to RNAi, we further validated these genetic interactions utilizing whole animal mutants for *dop* and *Apc*. Once again we assessed the number of brains with disrupted glomerular organization when one copy of *dop*, *apc*, or *fat2* were disrupted and compared these heterozygotes with transheterozygotes where we halved *fat2* expression by introducing one copy of *fat2* null allele (*fat2^N^*^103–2^) into *dop* or *Apc* heterozygotes. *dop* heterozygotes brains presented with few disruptions comparable to our controls (11% disrupted brains, 4/35 brains). When we added one copy of *fat2* null allele, we found a trending increase in the percentage of disrupted brains (21%, 7/33 brains, p=.33) (Fig 8C-D). The *dop*^1^ allele is known to be a hypomorph which produces the wildtype protein with a mutation in *dop* kinase domain leading to lowered/disrupted kinase activity ^58^. This incomplete disruption of the *dop* protein could mask possible genetic interaction between *fat2* and *dop*.

Therefore, further investigations are necessary to assess the trending increase in phenotypic penetrance for *dop*^1^*-fat2^N^*^103–2^ transheterozygotes. *Apc* and *Apc2* have been shown to play both mutually exclusive roles as well as redundant and synergistic roles ^65^. Therefore, to account for compensation from *Apc2* we used a whole animal with null mutations in both Apc family members in order to test genetic interaction with Fat2. Mutants with one copy of *Apc^Q^*^8^ and *Apc2^N^*^175^*^K^*null alleles had a significantly higher rate of disrupted brains compared to control (32%, 12/37 brains). Unlike *dop*^1^*-fat2^N^*^103–2^ transheterozygotes, *Apc^Q^*^8^*Apc2^N^*^175^*^K^* with one copy of the *fat2* null allele was able to significantly rescue the phenotype seen in *Apc^Q^*^8^*Apc2^N^*^175^*^K^* heterozygotes (11%, 4/35 brains). Given that *Apc* and *Apc2* encode proteins that play roles in multiple cell biological processes, it is unclear if Fat2 regulates Apc/Apc2’s role in the regulation of β-catenin/Wingless signaling or the regulation of microtubule organization. The bidirectional impact of Fat2, suggests that Fat2 likely balances cytoskeletal regulators to achieve the desired ORN class-specific axon behavior. It is possible that each ORN harbors different cytoskeletal regulators, and differential interactions of these components with Fat2 may explain class-specific effects of Fat2 and cytoskeletal regulators to orchestrate class-specific axon behaviors.

## Discussion

The olfactory sensory circuit possesses a distinctive feature present in nearly all neural circuits across the animal kingdom, namely its functionally organized topographic map. This map relies on widely dispersed neurons of the same type, which converge their axons into a neuropil called a glomerulus. The intricate process of how neuronal identity influences circuit organization is a significant area of research in neurobiology due to its close connection to neurodegeneration and neuronal dysfunction. Within the olfactory system, various cell surface proteins, such as Robo/Slit and Toll receptors, regulate many aspects of circuit organization, including axon guidance and synaptic matching. In this study, we have discovered an unconventional Cadherin protein called Fat2, which acts as a regulator of axon organization specific to certain classes. Fat2 exhibits the highest expression levels during metamorphosis, when the larval olfactory system disintegrates before re-constructing the mature olfactory circuit, with varying levels of Fat2 expression specific to each class. By studying *fat2* null mutants, we observed distinct effects on each class’s glomerulus, with the most severely affected being those with the highest expression of fat2. Furthermore, we provide evidence suggesting that the intracellular domain of Fat2 is necessary for its function in organizing the axons of olfactory receptor neurons (ORNs). Examination of confocal images from early pupal development indicates that Fat2 is crucial for appropriate axon retraction, which in turn allows for the condensation of class-specific neuropils. Our research also indicates that the expression of Fat2 in projection neurons (PNs) and local interneurons (LNs) does not significantly contribute to the organization of ORNs. Finally, we have identified potential interactors of Fat2’s intracellular domain that likely play a role in coordinating the necessary cytoskeletal changes for axon retraction during the early stages of glomerular development. In summary, our findings lay the groundwork for understanding the role of Fat2 in the organization of the olfactory circuit and highlight the critical importance of axon retraction during the maturation of protoglomeruli.

### Fat2 function in mediating ORN axon-axon interactions

ORN axons arrive at the antennal lobes early in pupal development and undergo a sequence of processes: defasiculation and dispersal across AL, class-specific axon compaction to form protoglomeruli, followed by an intra-glomerular axon expansion, and finally secondary compaction of the axon terminals as glomeruli form and mature (Fig 5). These axonal behaviors are accompanied by cell surface protein mediated recognition of ORN-PN-LN terminals and dendrites to allow for proper synapse formation across these varying components of the antennal lobe glomeruli^43^. The developmental analysis of *fat2* mutant phenotype revealed that the major process that seems to be defective is in the final stage of glomerular formation where axon terminals are consolidated into defined compact glomeruli.

This phenotype differs from the function of N-Cadherin^4^, which mostly impacts the initial elaboration of axon terminals within glomeruli, suggesting sequential utilization of different Cadherins in the stepwise progression of glomerular organization in the antennal lobes.

The pattern of Fat2 expression in the developing Drosophila brain shows very specific localization largely to ORNs and a subset of LNs, but not PNs, likely indicating adhesive interaction between ORNs and LNs within glomeruli. However, cell type-specific RNAi knockdowns of *fat2* are associated with glomerular defects only when Fat2 function is disrupted in ORNs but not in LNs. In whole animal *fat2* mutants that exhibit glomerular defects, ORNs still retain their normal PN synaptic targets. These results suggest that Fat2 mainly functions to mediate ORN-ORN interactions rather than establishing connection specificity with other neuronal components of olfactory circuit organization within antennal lobe glomeruli.

### Cytoskeletal remodeling through Fat2 intracellular signaling function is conserved between flies and mammals

Axonal growth cones in developing neurons are highly dynamic and require tight regulation of the cytoskeleton to respond to adhesive factors driving axon-axon and axon-dendrite interactions during axon fasciculation and synapse formation, respectively. Utilizing mass spectrometry, several publications have reported the identification of cytoskeletal proteins that interact with the intracellular domain of mammalian orthologue of the Drosophila Fat2 and Fat3 to regulate neuronal projection behaviors ^36, 48^. In our genetic analyses, we found many of the putative intracellular interactors. Particularly, RNAi knockdown axonal phenotypes of *dop,* which regulates cytoskeletal organization and membrane growth, were enhanced by reducing the dose of *fat2*. In contrast, whole animal mutant analysis of *Apc/Apc2,* which encodes a β-catenin regulators/cytoskeletal anchors, showed the phenotype was suppressed by reducing the dose of *fat2*. These results implicate cell surface molecule mediated regulation of the cytoskeleton dynamics in proper ORN axon terminal organization. Compellingly, disruptions in the mammalian Fat3 intracellular domain are sufficient to generate an ectopic plexiform layer in the retina that resembles the ORN class-specific glomerular splits we see in the antennal lobes of *fat2* mutants ^66^. Further studies revealed that the ectopic plexiform layer was a result of defects in neurite retraction similar to the developmental defects we see with *fat2* mutants ^36^. In addition, the aberrant neurites failed to form mature synapses, suggesting defects in axon behaviors are accompanied by defects in synapse formation ^48^. This finding provides an explanation for the ORN neurodegeneration seen in our *fat2* mutants since it is well accepted that disruptions to synaptogenesis/synaptic function can lead to regressive processes such as neurodegeneration^67^. Further studies are required to confirm whether ORN synaptic number is affected in *fat2* null animals. Even though no reports of the role of Fat3 function in mammalian olfactory circuit development, multiple sources confirm the high expression of Fat3 in the developing mouse olfactory bulb ^29, 33, 34^, suggesting Fat3 is functionally relevant in the mammalian olfactory circuit development. Our study highlights the conserved function of mammalian Fat3 and its Drosophila orthologue Fat2, in neural circuit development and architecture.

### Fat2 function in axon organization versus planar polarity

Previously, Fat2 function in Drosophila was predominantly proposed to be restricted to the regulation of planar cell polarity and collective cell migration ^23, 25, 68, 69^. In this paper, we show that Fat2 function in neuronal development and ORN axon terminal organization depends on the intracellular domain, as *fat2* mutants with only the extracellular domain phenocopy null mutants. This contrasts with the planar cell polarity phenotypes in the egg chambers where the extracellular domain of Fat2 alone can partially rescue the mutant phenotypes observed in *fat2* null mutants. As an atypical Cadherin, Fat2 can mediate cell-cell adhesion, yet membrane proteins that act as ligands or receptors for Fat2 have not been identified. Recent studies have implicated putative roles for Sema5c or Lar as potential trans interactors of Fat2 in mediating planar polarity and collective cell migration in the egg chambers ^68^; however, we were unable to detect genetic interactions in glomerular organization using mutations in either gene (data not shown). In addition, mutations in *Lar* exhibited phenotypes that were distinct from *fat2* mutants (data not shown). Therefore, these results suggest potential differences in mechanisms of Fat2 function in establishing planar cell polarity versus glomerular organization.

In addition to Fat2, another member of the family, Drosophila Fat cadherin, is a well-established regulator of growth and patterning via the Hippo pathway ^32, 70^. Despite their functional similarity, the intracellular domain of Fat and Fat2 are highly divergent and are much more similar to their mammalian orthologues, Fat4 and Fat3 respectively ^33, 71^. Furthermore, mammalian Fat1/Fat4 interaction can regulate aspects of neurodevelopment in mice ^72^. Thus, the function of Fat protein family members in supporting neuronal survival might be conserved across protein family members and species, as *fat* mutants are associated with ommatidial degeneration in the Drosophila visual system ^73^. It is unclear if Fat protein function in the nervous system extends outside of the visual system or interacts with Fat2 in axon terminal organization. Another future avenue to investigate is whether interactions between Fat2 and Fat, or other transmembrane receptors, drive olfactory circuit organization and neuronal survival.

### Limitations

Our analysis of *fat2* expression profile suggests high expression in the local interneurons ^1^ during olfactory circuit development. Yet, LN-specific knockdown of *fat2* did not impact ORN axon organization. Given that the scope of our research focuses on ORN class-specific regulation of ORN circuit topography, the role of Fat2 in LN development remains to be investigated. LNs play an integral role in modulating synaptic sensitivity to appropriately relay inter-glomerular and intra-glomerular signals^74^. Fat2 regulation of glomeruli morphology and expression in LNs, a key regulator of olfactory signal processing, raises the fundamental question as to whether *fat2* null flies exhibit behavioral defects. Behavioral analysis and electrophysiological evaluations of *fat2* null/ICD mutants or cell type-specific knockdown of *fat2* could elucidate possible neuro-pathologies related to *fat2* dysfunction. This could provide possible mechanistic insights into mutations in mammalian *fat3* associated with schizophrenia and bipolar disorder^75^.

While we have shown that the extracellular domain of Fat2 is not sufficient to rescue the fractured class-specific glomeruli, the extracellular domain of many cadherin proteins remains integral to their function, and Fat2 is likely no exception^20^. To directly investigate the role of Fat2 extracellular domain (*ECD*), we would need to generate a *fat2^ΔECD^* mutant with a functional intracellular domain and disrupted extracellular domain. However, due to the size of the extracellular domain, molecular cloning techniques to precisely remove the ECD have not been successful. Similarly, the generation of an insertion mutant, *UAS-fat2,* is difficult due to the sheer size of the *fat2* gene. A *GAL4* inducible *UAS-fat2* would be a powerful reagent because it would allow us to perform rescue experiments as well as over-expression analysis. Recent advances in gRNA-based overexpression using modified transcriptional activator CRISPR proteins can also be useful approaches for rescuing mutant phenotypes and overexpression experiments further probing Fat2 function.

Given that *fat2* likely functions during dynamic axon extension/retraction, available tools for inducible genetic disruptions, including temporally specific or temperature-sensitive genetic drivers, may not be precise enough to assess *fat2* function. Furthermore, developmental trajectories of ORN classes are highly variable and dependent on when a given ORN class is born^6^, which can lead adjacent glomeruli to be at very different developmental phases at the same time point. Therefore, knocking down *fat2* at a specific time point will affect different axon behaviors depending on the developmental phase of each ORN class. It is yet unclear whether the expression of *fat2* in adjacent glomeruli can affect each other’s glomerular organization, and thus we cannot directly associate resulting phenotypes with *fat2*’s role in axon extension/retraction.

A major limitation of our study is the absence of live single-axon imaging. Our developmental analysis provides snapshots of axonal bundles to investigate general disruptions in axon behavior in *fat2* null mutants (Fig 5). This coarse perspective may obscure potential roles that *fat2* plays in more transient axon behaviors, as well as *fat2*-dependent variations in behavior between single axons in the same ORN class. Additionally, live imaging of Fat2 protein dynamics in conjunction with cytoskeletal markers or intracellular interactors identified in this paper could further explain how Fat2 can both enhance and repress glomerular disruptions. Our paper identifies the discrete developmental time point and the cellular expression profile for further investigations into Fat2 function in olfactory circuit organization.

### Future directions and conclusion

To summarize, one general organizational logic found in neural systems across various species is the arrangement of neurons performing similar functions to project to similar regions of the brain. Our study provides a novel candidate regulator, Fat2, of this neuronal class-specific organizational principle and suggests there may be an intracellular combinatorial code of cytoskeletal effectors, in conjunction with the cell surface combinatorial code. Further investigation will be necessary to identify the mechanism behind Fat2-APC/APC2 interactions as well as validate Fat2-Dop interactions. Using these intracellular combinatorial codes, we can begin to leverage drugs that target the intersection of intracellular effectors and cell surface molecules to enhance the precision of pharmacological interventions. While investigating Fat2 function, we also found evidence of sequential class-specific axon convergence events during glomerular development (Fig. 5). This finding provides a more stepwise understanding of olfactory system development, and hints that each axon convergence step is mediated by unique sets of cell surface proteins. Given that Fat2 protein is consistently present throughout olfactory system development, what mechanisms prevent Fat2 from regulating axon convergence in other axon convergence steps? Also, can Fat2 regulate axon convergence outside of a developmental context, such as during regeneration? Finally, our results provide evidence for a conserved Fat2 function in organizing both the mammalian visual system and insect olfactory circuit architecture, thus emphasizing the translational implications^76^ of further research into Fat2 and the Fat cadherin family.

## Methods

### Key Resources Table

Genetic reagents
| Figure/Experiment | Name | Source/Reference | Identifier/BDSC # |
| --- | --- | --- | --- |
| Fig 4 | *fat2ΔICD-3xGFP FRT80B* | Laboratory of S. Horne-Badovinac; PMID:28292425 |  |
| Fig 1 and 2 | *fat2N103-2 FRT80B* | Laboratory of Sally Horne-Badovinac; PMID:22413091 |  |
| Fig 1 | *Or47b-CD2/CYO; TM2/TM6B* | Leslie Vosshall PMID: 16139209 |  |
| Fig 1 | *UAS-fat2 RNAi (kug)* | FBgn0261574 | BDSC 40888 |
| Fig 1 and 4 | *Or47b-GAL4 Or23a-GAL4 Or47a-Gal4 Gr21a-GAL4 UAS-SytGFP* | recombined from Vosshall et al. 2000; Fishilevich and Vosshall 2005 |  |
| Fig 1 | *fat2-58D* | PMID: 19906848 |  |
| Fig 2 | *Fat2-3xGFP FRT80B* | Laboratory of S. Horne-Badovinac; PMID:28292425 |  |
| Fig 2 | *UAS-mCD8GFP* | Laboratory of P. Volkan |  |
| Fig 2 | *eyFLP/+;; UAS>STOP>mCD8::GFP/+* |  | BDSC 30032 |
| Fig 2 | *Fat2-GAL4 (kug)* |  | BDSC 77610 |
| Fig 3 | *Or19a-GAL4* |  | BDSC: 24617 |
| Fig 3 and 4 | *Or69a-GAL4* | PMID:21940430 | BDSC 9999 |
| Fig 3 | *Ir40a-GAL4* | PMID:21940430 | BDSC 41727 |
| Fig 3 and 4 | *Ir64a-GAL4* | PMID:21940430 | BDSC 41732 |
| Fig 3 and 4 | *Ir84a-GAL4* | PMID:21940430 | BDSC 41734 |
| Fig 3 | *Or22-GAL4* | PMID:21940430 |  |
| Fig 3 | *Or88a-mCD8GFP* | Leslie Vosshall PMID: 16139209 |  |
| Fig 5 | *72OK-GAL4 UAS-sytGFP* | Laboratory of P. Volkan |  |
| Fig 3 | *UAS-sytGFP/CYO; TM2/TM6B* | Laboratory of P. Volkan |  |
| Fig 4 | *10xUAS-mCD8RFP* |  | BDSC 32219 |
| Fig 6 | *(orb) 499-GAL4* |  | BDSC 63325 |
| Fig 7 | *peb-GAL4; Or47b/Or23a/Gr21a-SytGFP/CYO; TM2/TM6B* |  |  |
| Fig 7 | *Apc* |  | BDSC 34869 |
| Fig 7 | *CG17528 (zyg8)* |  | BDSC 42575 |
| Fig 7 | *Chb* |  | BDSC 35442 |
| Fig 7 | *dop* |  | BDSC 60138 |
| Fig 7 | *dop1 red1 e1/TM6B, Tb1 (dop1)* |  | BDSC 64424 |
| Fig 7 | *Apc2N175K ApcQ8/TM3* |  | BDSC 7211 |

#### Genetic reagents

##### Antibodies

| Name | Company | Cat. # | Conc used |
| --- | --- | --- | --- |
| Goat anti-Rabbit IgG (H+L) Highly absorbed secondary antibody, Alexa Fluor Plus 488 | Invitrogen | Ref:A32731<br>Lot:XH353659 | 1:1000 |
| Alexa Fluor 546 goat anti-mouse IgG (H+L) | Invitrogen | Ref:A11030<br>Lot:2273717 | 1:300 |
| Anti-green fluorescent protein, rabbit IgG fraction | Invitrogen | Ref:A11122<br>Lot:2477546 | 1:1000 |
| Rat anti-NCAD (DN-Ex #8) | DSHB |  | 1:20 |
| Mouse anti-rat CD2 | BioRAD | MCA154R Batch<br>No:1611 | 1:500 |
| Alexa Fluor 647 Goat anti-rat IgG (H+L) | Invitrogen | Ref: A21247<br>Lot:2311802 | 1:200 |

##### Genetics, fly husbandry, and RNAi knockdown

Flies were kept at room temperature or 25 degrees. For RNAi experiments, RNAi crosses were reared at 28°C and only moved to RT (room temperature) on the day of dissection. Flies were dissected 3-7 days after eclosion. To restrict background variation, we collected RNAi knock down only, No GAL4 control, and genetic interaction flies from the same cross.

ORN specific knockdown of *fat2* RNAi utilized *pebbled-GAL4; UAS-fat2 RNAi* (BDSC:40888); Or47b-mCD8GFP. LN specific knockdown of *fat2* RNAi utilized *UAS-kug RNAi* (BDSC:40888); Or47b-mCD8GFP/ *449-GAL4* (LN-subtype specific marker, BDSC:63325).

##### Fat2 protein expression analysis and intersectional labeling

Developmental analysis of Fat2 protein expression was obtained using CRISPR generated Fat2-3xGFP flies graciously gifted to us by the Horne-Badovinac Lab (Horne-Badovinac et al., 2012). White pupa (staged as 0 APF) were collected on a microscope slide, then placed in a closed petri dish with a damp kim wipe for ∼24hrs, ∼48hrs, and ∼72hrs (may vary ±3 hours). Both brains and antenna were dissected, immunostained, and imaged.

To intersectionally label *fat2* expressing cells we used cell type specific FLP/FRT (*ey-FLP* for ORN specific labeling, and *GH146-FLP* for PN specific labeling) in conjunction with *fat2-*GAL4 (BDSC:77610) and *UAS>stop>GFP* (BDSC: 30032) system to intersectional label cell type specific expression patterns for *fat2.* Time point collection, dissection, and imaging performed according to immunohistochemistry methods section.

##### Immunohistochemistry, Image Acquisition, and Statistics

Flies were immobilized by and washed in 70% ethanol for 30 seconds and then moved to .1% PBT (Phosphate Buffer Solution with .1% Triton X-100). Brains were removed via micro dissection and immediately placed in .4% PFA (paraformaldehyde) on ice. Then the brains were fixed in 4% PFA for 15 to 30 minutes at room temperature. Then washed in PBT 3x for 20 minutes each wash.

For immunostaining primary antibodies were used at the following concentrations: rabbit α-GFP 1:1000 (Invitrogen), rat α-Ncad 1:20 (Developmental Studies Hybridoma Bank), mouse α-rat CD2 1:200 (Serotec),

Secondary antibodies were used at the following concentrations: AlexaFluor488 goat α-rabbit 1:1000, AlexaFluor568 goat α-mouse IgG highly cross-adsorbed 1:300, AlexaFluor647 goat α-rat 1:200. Confocal images taken using an Olympus Fluoview FV1000 at 40x.

*P*-value was calculated by two-tailed Fisher’s exact test through the built-in functions of GraphPad Prism 9 software.

Stacks of optical sections 2 μm thick were collected on Olympus Fluoview FV1000 of the entire antennal lobe. Z-stack projections created on either Fluoview or FIJI and then imported into Adobe Illustrator to generate figures.

##### 72OK-GAL4 Developmental time point analysis

One batch of brains between 18 hAPF and 30 hAPF were dissected, imaged, and then visually assessed for age^3, 4, 39^. Another batch of brains 24 hAPF to 50 hAPF were dissected, imaged, and then visually assessed for age. Developmental time point was determined using brain morphology found in both groups indicate brains aged between 24 hAPF to 30 hAPF.

##### Analyzing the *fat2* expression in the single-cell RNA-seq datasets

The annotated single-cell RNA-seq datasets were from McLaughlin elife 2021 (GSE162121). We extracted the expression values (log_2_(CPM +1)) of *fat2* (*kug*) across all the cells assigned to ORN classes. We calculated the fraction of *fat2*-positive cells (defined by log_2_(CPM +1) ≥ 1) and the mean expression of *fat2* of each ORN class at each developmental stage.

## Data availability

All relevant data are within the paper and its supporting information files.

## Acknowledgements

We are grateful to Sally Horne-Badovinac for sharing Fat2 genetic reagents (as well as *sema5c* and *lar* mutants), and Christian Dahmann for sharing *fat2-58D* and other *fat2-related* fly lines. We want to thank Rachel Estrella, Douglas Blackiston, Julie Reynolds, Kavya Raghunathan, Latanya Coke, and Volkan lab members for their input on the manuscript. We thank the Bloomington Stock Center for its services.

## Funding

This study was supported by National Science Foundation award 2006471 to PCV and the National Science Foundation Graduate Research Fellowship under Grant No. DGE 2139754 to KMV.

## Conflicts of interest

The authors declare that they have no competing interests.

## Author contribution

Conceptualization: KMV and PCV. Investigation: KMV and CY. Analysis/interpretation of data: KMV, QD, and PCV. Manuscript-original draft: KMV, QD, PCV. Manuscript-revision: KMV and PCV.

## Supplemental Figures 1-5

**Supplemental Figure 1.**
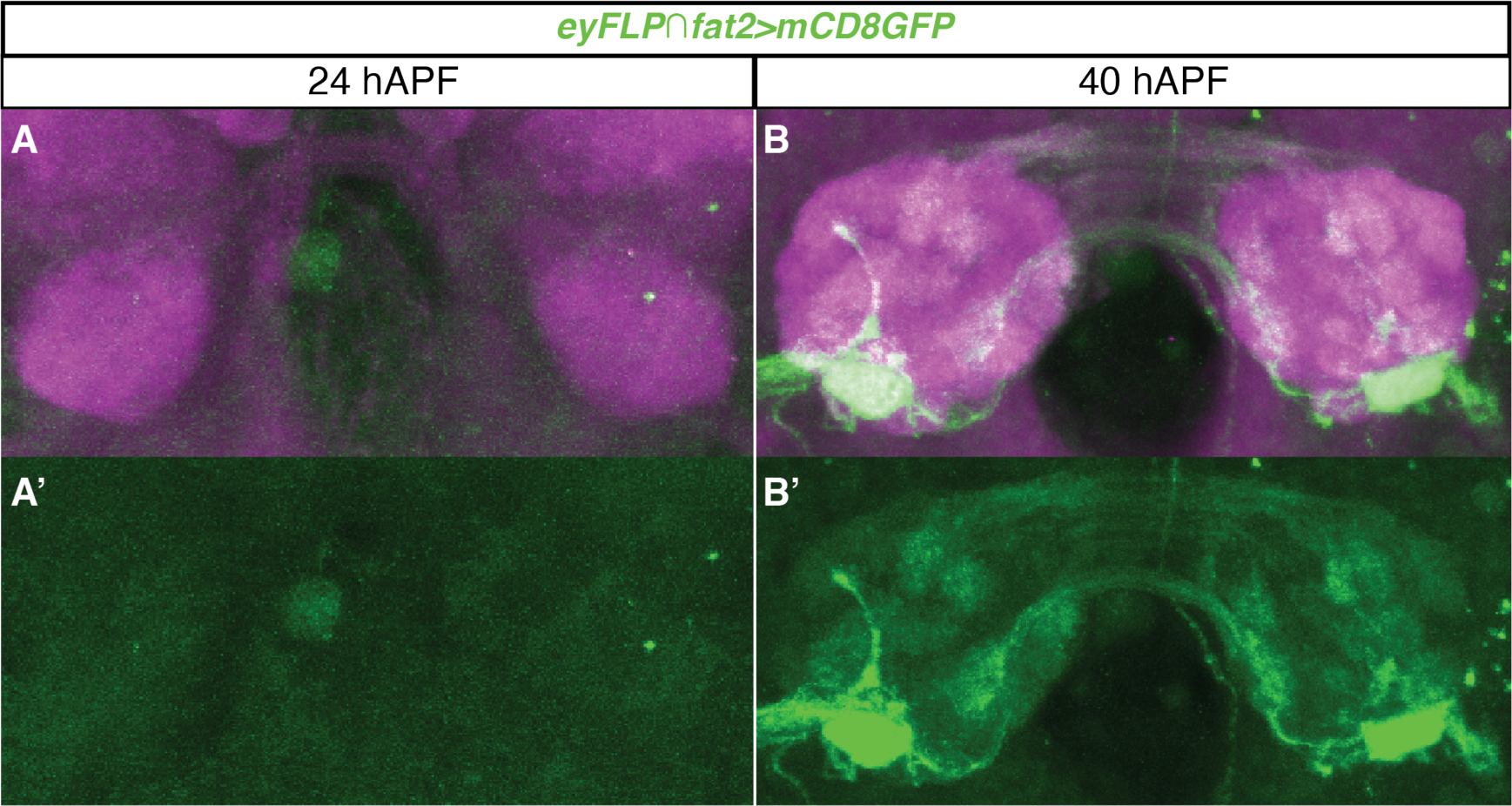
ORN expression of *fat2* starts between 24 hAPF and 40 hAPF. **(A-B’)** Confocal images of *eyFLP*, *fat2-GAL4, UAS<STOP<mCD8GFP* antennal lobes at 24 hAPF and 40 hAPF. Little to no GFP signal present in the antennal lobes at 24 hAPF.

**Supplemental Figure 2.**
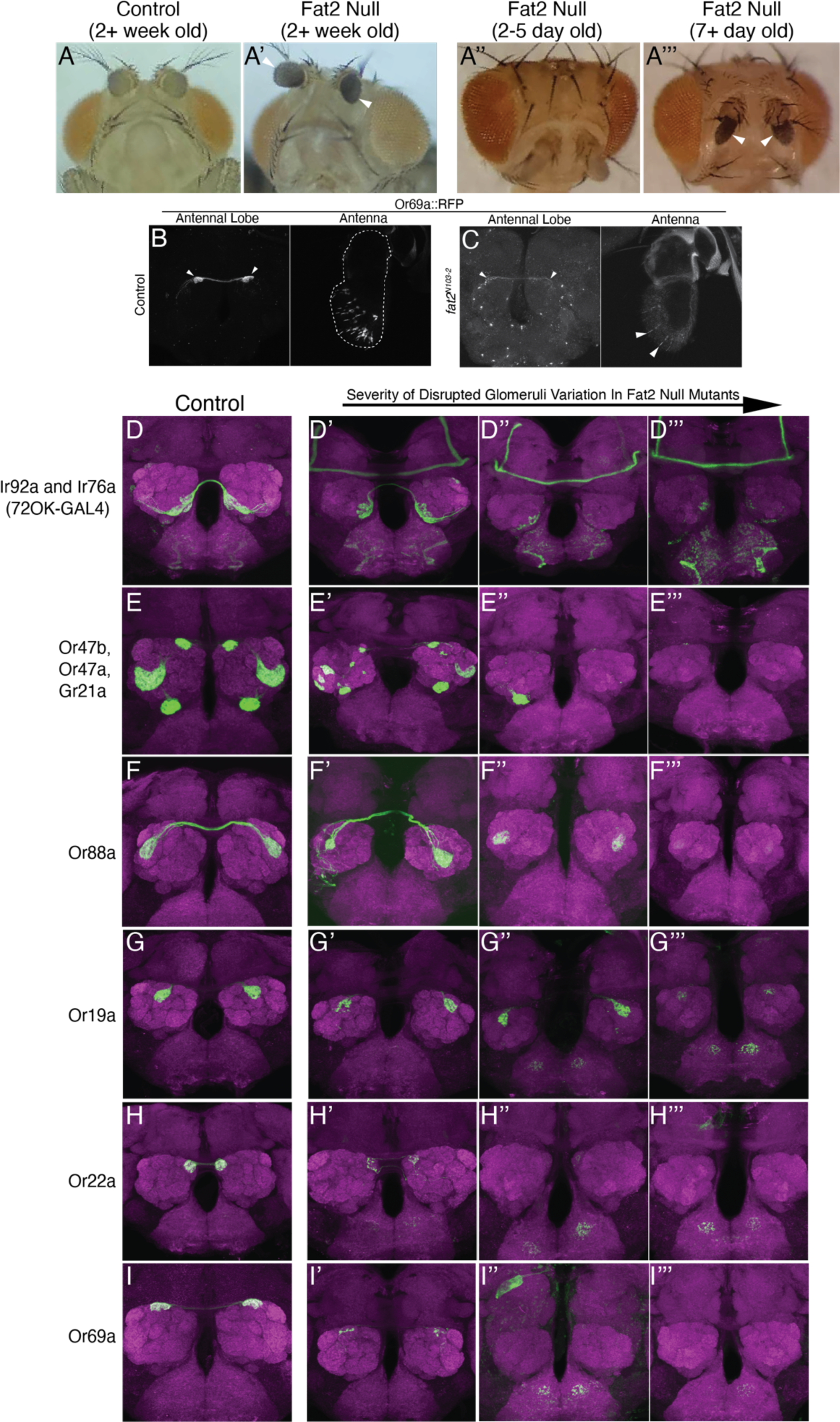
Sensory neuron degeneration in *fat2 null* mutants. **(A and A’)** Brightfield images of live, anesthetized, two weeks old age-matched control and mutant (homozygous *fat2^N^*^103–2^) siblings from the same vial. White arrows point to discoloration found on mutant fly antenna. Darkening of tissue often indicative of cellular damage. **(A’’-A’’’)** Brightfield images of homozygous *fat2* null mutants at two different ages. White arrows indicate the appearance of discolored antenna on older mutant flies but not younger mutant flies. **(B-C)** Confocal images of antennal lobe and antenna from both control and *fat2 null* mutant flies labeled with *Or69a-GAL4 10xUAS-RFP*. Dotted line outlines the second and third antennal segments. White arrows point to the Or69a cell bodies in mutant antenna and Or69a axon terminals in the antennal lobe. Background signal higher in mutant brains and antennal images because images were taken with higher laser intensity to visualize the few fluorescent cells present. **(D-I)** Control brains labeled with different *OrX-GAL4* drivers to show the respective glomeruli shape and position. **(D’-I’’’)** Representative images of brains with increasing severity of disruption to glomerular shape and position, with the most severe disruption being the absence of glomerular signal altogether in the last column. Note the lack of clear glomerular boundaries in the last column. The classes in this figure were selected because they displayed the absence of glomerular fluorescence.

**Supplemental Figure 3.**
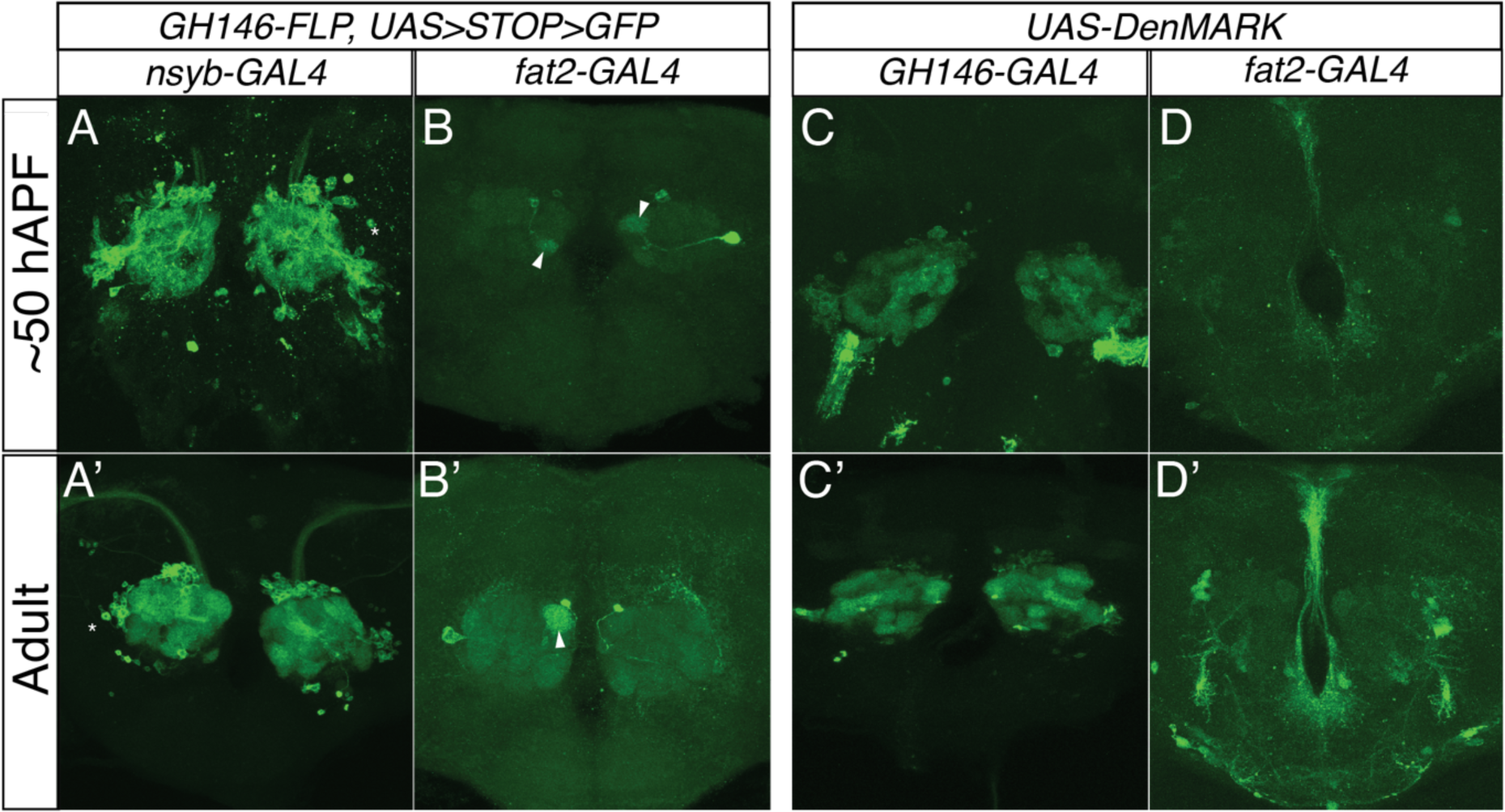
PNs express very low to undetectable levels of *fat2*. **(A and A’)** Confocal images of adult and 50 hAPF pupal antennal lobes from *GH146-FLP*, *nsyb-GAL4, UAS<STOP<GFP*, which labels *GH146-*labeled PNs that also express *nsyb*. **(B and B’)** Adult and mid-pupal antennal lobe images from *GH146-FLP*, *fat2-GAL4, UAS<STOP<GFP* which labels *GH146-*labeled PNs that also express *fat2.* **(C and D’)** Adult and mid-pupal antennal lobe images labeling dendritic projections from either *GH146-*labeled neurons or *fat2+* neurons.

**Supplemental Figure 4.**
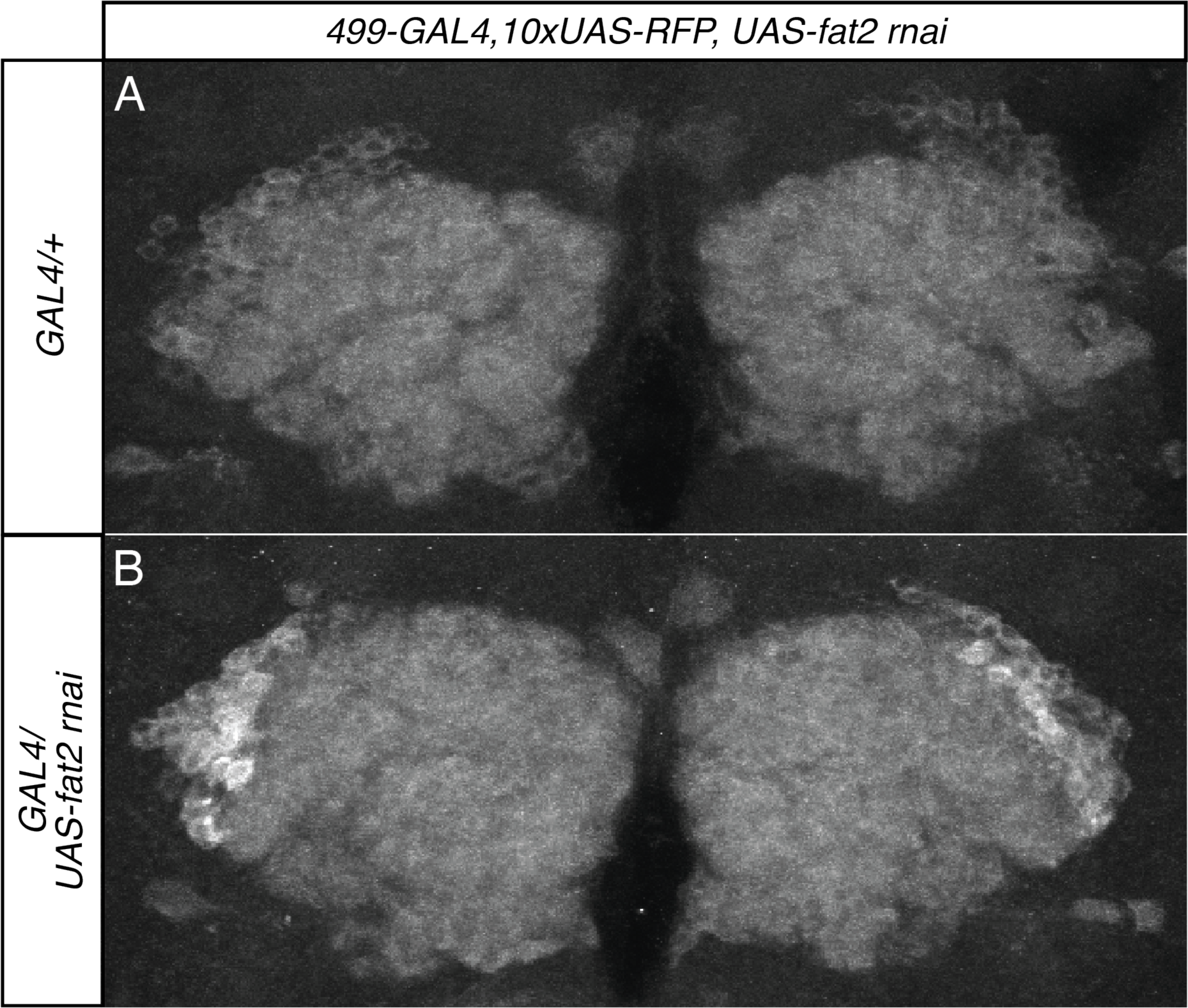
o*r*b*+(499-GAL4)* LNs neurite terminal organization display no discernible aberrations upon *fat2 RNAi* knockdown. **(A and B)** Confocal images of adult antennal lobes from *499-GAL4, 10xUAS-RFP*, which labels *orb+* LN cell bodies and projections. **(A)** shows LN projection pattern for *UAS-fat2 RNA*i control and **(B)** shows projection pattern when *fat2* is knocked down specifically in *orb+* LNs.

**Supplemental Figure 5.**
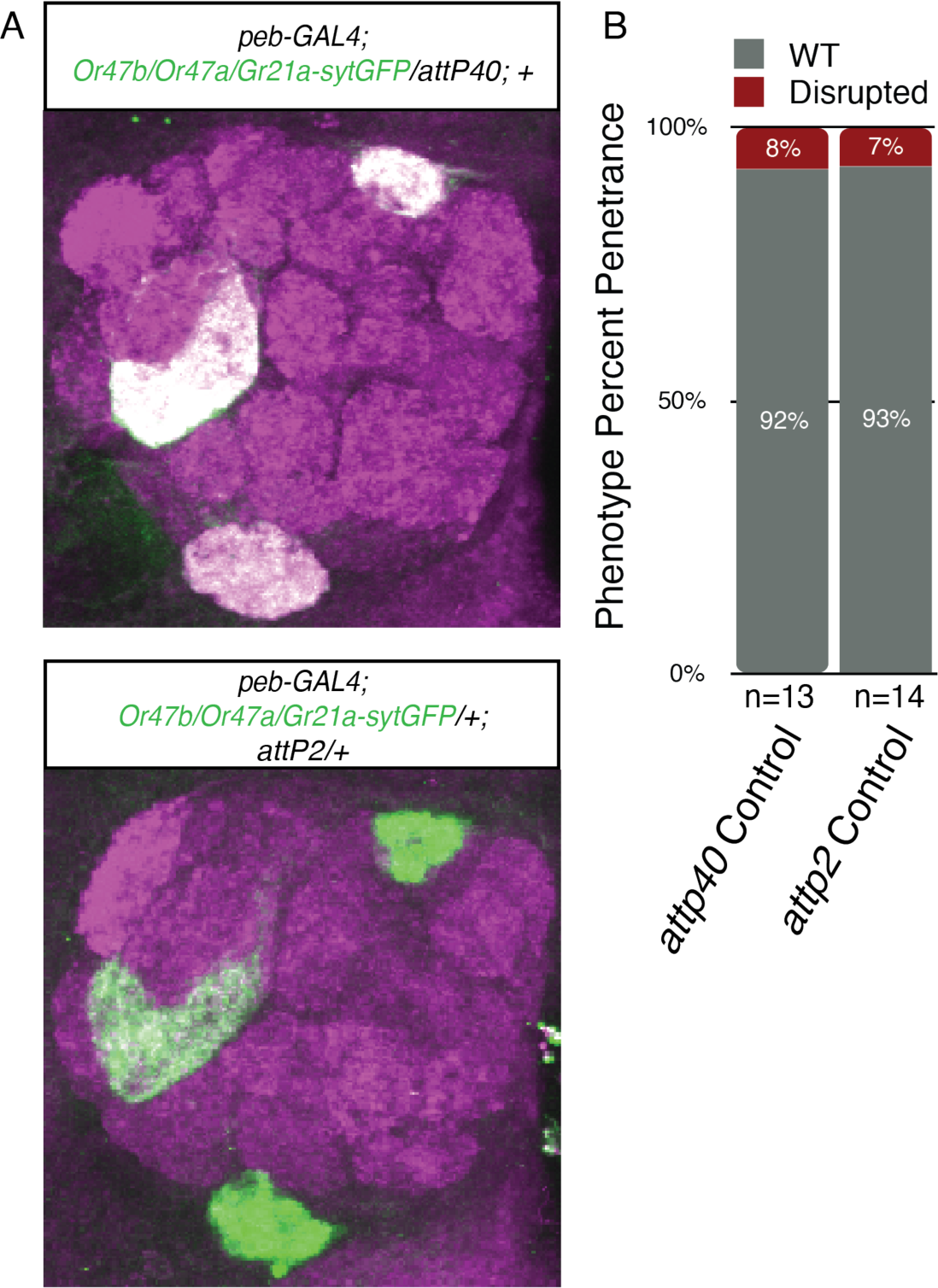
Heterozygous attp40 and attp2 empty insertion sites do not disrupt glomerular organization for glomeruli innervated by Or47a, Or47b, and Gr21a. **(A)** Representative confocal images of antennal lobes from the respective genotypes. **(B)** Bar graphs to visualize quantification of phenotypic penetrance for each respective genotype

## Notes

### Competing Interest Statement

The authors have declared no competing interest.

